# DNA Binding Induces a *cis* to *trans* Switch in Cre Recombinase to Enable Intasome Assembly

**DOI:** 10.1101/2020.05.24.113654

**Authors:** Aparna Unnikrishnan, Carlos D. Amero, Deepak Kumar Yadav, Kye Stachowski, Devante Potter, Mark P. Foster

**Author notes:** Corresponding Author: Mark P. Foster) Department of Chemistry and Biochemistry, Columbus, OH 43210. CLASSIFICATION: Biological Sciences: Biochemistry.

## Abstract

Mechanistic understanding of DNA recombination in the Cre*-loxP* system has largely been guided by crystallographic structures of tetrameric synaptic complexes. Those studies have suggested a role for protein conformational dynamics that has not been well characterized at the atomic level. We used solution NMR to discover the link between intrinsic flexibility and function in Cre recombinase. TROSY-NMR spectra show the N-terminal and C-terminal catalytic domains (Cre^NTD^, Cre^Cat^) to be structurally independent. Amide ^15^N relaxation measurements of the Cre^Cat^ domain reveal fast time scale dynamics in most regions that exhibit conformational differences in *active* and *inactive* Cre protomers in crystallographic tetramers. However, the C-terminal helix αN, implicated in assembly of synaptic complexes and regulation of DNA cleavage activity via *trans* protein-protein interactions, is unexpectedly rigid in free Cre. Chemical shift perturbations and intra- and inter-molecular paramagnetic relaxation enhancement (PRE) NMR data reveal an alternative auto-inhibitory conformation for the αN region of free Cre, wherein it packs *in cis* over the protein DNA binding surface and active site. Moreover, binding to *loxP* DNA induces a conformational change that dislodge the C-terminus, resulting in a *cis* to *trans* switch that is likely to enable protein-protein interactions required for assembly of recombinogenic Cre intasomes. These findings necessitate a re-examination of the mechanisms by which this widely-utilized gene editing tool selects target sites, avoids spurious DNA cleavage activity, and controls DNA recombination efficiency.

**SIGNIFICANCE STATEMENT:** The Cre-*loxP* system is a widely used gene editing tool that has enabled transformative advances in immunology, neuroscience and cardiovascular research. Still, off-target activities confound research results and present obstacles to biomedical applications. Overcoming those limitations requires understanding the steps leading to assembly of recombination complexes, *intasomes*. We measured the magnetic properties of nitrogen nuclei in the backbone of the enzyme to correlate its intrinsic dynamics with its function in DNA recognition and cleavage. Remarkably, we found that in the absence of DNA the C-terminus of Cre appears to block the DNA binding surface and active site of the enzyme. Binding to *loxP* DNA induces a conformational switch that would enable the intermolecular protein-protein interactions required for assembly of recombinogenic Cre intasomes.

## INTRODUCTION

Site specific DNA recombinases represent an attractive option for genome engineering – the insertion or exchange of genes into precise locations in chromosomes^1–4^. The tyrosine recombinase family of phage-derived enzymes (e.g., λ-integrase, Cre and Flp recombinases), which evolved to facilitate viral infection, gene transposition and bacterial pathogenesis, have proven useful for applications that include DNA subcloning without restriction enzymes, and conditional expression of target genes^5–7^. These proteins bind specifically to pairs of inverted short palindromic DNA sequences (*recombinase binding elements*; RBEs) and mediate recombination by assembly of tetrameric *intasomes* (Fig. 1*A* and *B*) which perform concerted DNA strand cleavage, exchange and ligation reactions^5,8,9^. Compared to other genome engineering approaches^10,11^, site specific DNA recombinases have the advantage that they allow a high degree of specificity, don’t require involvement of additional host-encoded factors, and are capable of generating cleanly integrated double-stranded DNA products^3,12–14^. These advantages motivate efforts to understand and manipulate the mechanisms of this family of enzymes.

**Figure 1.**
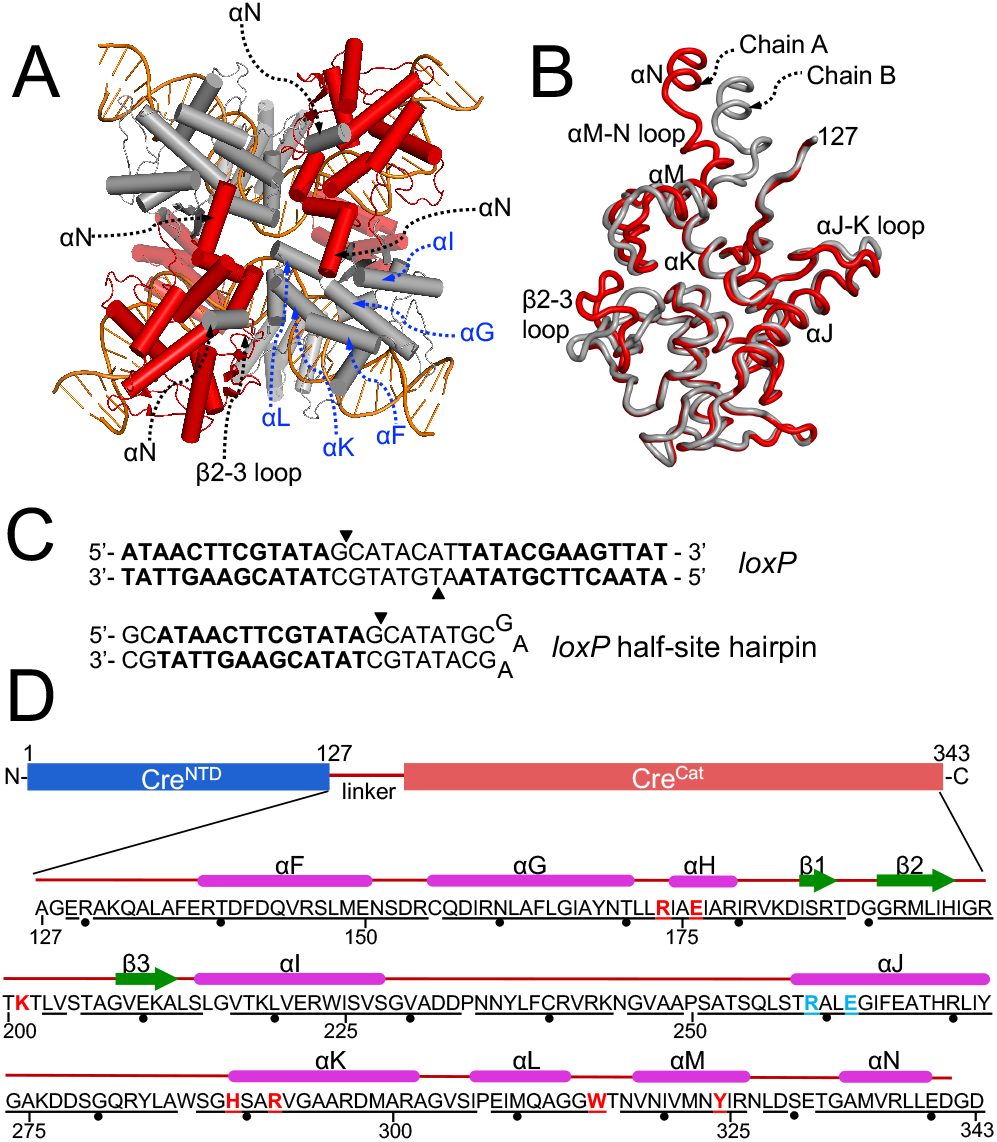
Conformational changes underlie coordinated DNA recombination by Cre recombinase. *(A)* Tetrameric synaptic complex of four Cre protomers bound to two *loxP* DNA duplexes (PDB ID 2HOI), viewed from the catalytic domain, Cre^Cat^. C-terminal helix αN of each protomer packs into a *trans* surface cavity (labeled in blue) in the adjacent protomer. *(B)* Superposition of Cre^Cat^ domains from “inactive” (red) and “active” (gray) Cre protomers (PDB ID 2HOI: chains A and B, respectively) highlight differences in the β2-3 loop, helix αM, αM-N loop and C-terminal helix αN. *(C) loxP* DNA, and the *loxP* half-site hairpin DNA sequence used in these studies; arrows indicate cleavage sites in full-length *loxP* DNA. *(D)* Domain map of Cre recombinase. Residues 1-126 (blue) comprise the N-terminal domain (Cre^NTD^). The interdomain linker and C-terminal catalytic domain comprise the Cre^Cat^ construct (residues 127- 343) used in these studies (pink). Sequence of the Cre^Cat^ construct is shown (numbered 127 to 343; dots shown to guide) with corresponding secondary structure elements from synaptic complex crystal structure (PDB ID 2HOI, chain B); residues G342 and D343 are not modeled in electron density maps. Active site and DNA binding specificity determinant residues are highlighted in red and cyan respectively. Backbone amide NMR chemical shift resonance assignments were made for all underlined residues.

Cre (<u>C</u>auses <u>R</u>ecombination) is the best studied member of the tyrosine recombinase family, yet fundamental gaps exist in our understanding of its mechanism of DNA site selection, intasome assembly, and allosteric control of DNA cleavage. The overall mechanism of Cre-mediated DNA recombination has been mapped out via many biochemical and crystallographic studies^5,15–19^. The enzyme catalyzes recombination between a pair of homologous 34 bp *loxP* sites (Fig. 1*C*) via a highly orchestrated series of events involving recognition and binding of a pair of Cre molecules to its semi-palindromic recognition site containing two RBEs (*loxP* half-sites), “synapsis” of an antiparallel pair of Cre dimers bound to *loxP* to form a stable tetrameric structure, coordinated cleavage of two opposing DNA strands to produce an intermediate with two of the Cre protomers covalently attached to a 3’-phosphate via a tyrosine linkage, followed by strand transfer and re-ligation to form a four-way DNA Holliday junction intermediate^5^. Two-fold asymmetry in the synaptic structures (Fig. 1*A*) is implicated in regulation of DNA strand cleavage activity, and thereby influence the order of strand cleavage and direction by which the Holliday junction is resolved^17,20,21^.

Although much is known about the overall mechanism of the reactions catalyzed by Cre and related site-specific recombinases, the nature of the conformational changes in the protein and protein-DNA complexes that facilitate the various steps in the pathway remain poorly understood^16,17,19,22–25^. Cre binds its target sequences by forming a C-shaped clamp with a C-terminal catalytic domain (Cre^Cat^) possessing the namesake tyrosine residue on one side of the DNA, and on the other an N-terminal domain (Cre^NTD^), connected by an extended 8 amino acid linker (Fig. 1). A series of sequential inter-protomer protein-protein interactions across the tetrameric synapse distinguish the structures of the active and inactive protomers. Within the catalytic domain, the tyrosine residue that serves as the nucleophile to catalyze phosphoryl transfer and formation of the covalent intermediate is located on the penultimate helix αM, while the αM-N loop and C-terminal helix αN from each protomer makes a *trans* contact (in a “clockwise” manner as viewed in Fig. 1*A*) to the neighboring protomer in the tetrameric complex. The β2-3 loop of each protomer abuts its own helix αM, and in the tetramer packs proximally to helix αM of the neighboring protomer, in the opposite direction compared to helix αN. Structural asymmetry in tetrameric structures is localized to these two regions of the protein (Fig. 1*A* and *B*). The asymmetry in the C-terminal and inter-protomer interfaces observed in the synaptic complexes of Cre and other tyrosine recombinase family members suggest that the intrinsic dynamic behavior of the enzymes is important in controlling its function, both in mediating tetramer assembly, and for regulating protomer activity^26–32^. Despite the importance of understanding the basis for conformational differences in Cre, structural insights have been largely limited to crystal structures of isosteric tetrameric synaptic complexes^5,16,17,19^.

We have used solution nuclear magnetic resonance (NMR) spectroscopy to explore the link between the intrinsic dynamic behavior of Cre and its function. First, NMR spectra of full-length Cre and of the isolated catalytic domain (Cre^Cat^) support the premise that the N-and C-terminal domains of Cre are structurally uncoupled. Nuclear spin relaxation experiments reveal flexibility in regions of Cre^Cat^ that are associated with both protein-protein and protein-DNA interactions. Unexpectedly, we found that the region of the protein comprising C-terminal helix αN is not highly dynamic, contrary to the expectation from crystal structures that it would be extended in solution^5,16,17,19^. Instead, NMR chemical shift perturbations and paramagnetic relaxation enhancement (PRE) experiments show the C-terminus of unbound Cre to be located in the DNA binding active site, in an apparent auto-inhibitory conformation. Upon binding to *loxP* DNA, the data show that the C-terminus is displaced, after which it is able to participate in fledgling intermolecular protein-protein interactions with other Cre molecules. These findings shed light on the previous paradoxical reports of *trans* interactions regulating Cre activity^33^, and call for a revised view of the reaction mechanism, highlighting the role of protein dynamics in regulating conservative site-specific DNA recombination.

## RESULTS

### NMR spin relaxation shows unexpected rigidity in the Cre C-terminal helix αN

To explore the role of protein dynamics in enabling the interconversion of Cre reaction intermediates in solution, we expressed and purified full-length Cre (38.5 kDa, residues 1-343) and a C-terminal fragment encoding the interdomain linker along with the C-terminal domain (23.8 kDa, residues 127-343). This domain, herein termed Cre^Cat^ (Fig. 1*B* and *D*) includes all seven active site residues (R173, E176, K201, H289, R292, W315 and Y324), the Cre-DNA binding specificity determinants (R259 and E262) as well as the C-terminal region implicated in *loxP* DNA binding cooperativity and synapsis (Fig. 1*A*)^17,33–37^. Both full-length Cre and Cre^Cat^ are predominantly monomeric in the absence of DNA, as indicated by their elution times in size exclusion chromatography (Fig. S1 *A*). Comparison of 2D ^1^H-^15^N TROSY-HSQC correlation spectra of Cre^Cat^ and full-length Cre showed that the well-dispersed signals overlay well (Fig. S1 *B*). This indicates that Cre^Cat^ folds independently in solution and that the N-terminal domain of Cre does not interact with the catalytic domain in solution, thereby justifying structural and dynamics studies of the isolated domain.

For backbone resonance assignments of Cre^Cat^ we expressed, purified the [U-^15^N] and [U-^15^N,^13^C] labeled protein and recorded HSQC- and TROSY-based double- and triple-resonance NMR spectra. Homogeneity and integrity of protein constructs were verified by SDS-PAGE and mass spectrometry (data not shown). We obtained backbone resonance assignments for 195 of 214 non-proline amides (> 91%) (Fig. 1*D*, S2 and S3). Unassigned amide resonances include the two N-terminal residues (A127 and G128), and a few residues in the loop regions of Cre^Cat^, due to line broadening from intermediate conformational exchange.

To probe the fast timescale backbone dynamics in Cre^Cat^, we measured ^15^N NMR (R_1_, R_2_) and {^1^H}-^15^N heteronuclear NOE (HetNOE) relaxation data (Fig. 2). The data show that Cre^Cat^ backbone amides largely exhibits uniform relaxation rates, with an overall rotational correlation time (τ_*C*_) of 13.2 ± 0.9 ns, computed from the trimmed-mean R_2_/R_1_ ratios^38^. Distinctly lower R_2_/R_1_ ratios and decreased {^1^H}-^15^N HetNOE values indicating fast (ps-ns) internal motions, were observed for the linker residues through A134 preceding αF, the β2-3 loop, αJ-K loop, αM-N loop and a few residues after the C-terminal helix αN. The β2-3 loop forms inter-protomer contacts in Cre-DNA synaptic tetramers and adopts distinct conformations between the catalytically *active* and *inactive* forms of Cre (Fig. 1*A* and *B*)^5^. The data show the αJ-K loop to be flexible in free Cre^Cat^ though it does not differ structurally between the two Cre isomers, consistent with a role in DNA binding, not in regulating recombination^5^. Surprisingly, although residues bracketing the C-terminal helix αN exhibit evidence of fast internal motions, the relaxation rates for the helix αN (A334 - L339) are close to the average values, indicating that it tumbles with the catalytic core of the protein. This suggests that instead of being extended in solution as might be expected from the *trans* conformations observed in tetrameric crystal structures, αN might adopt a distinct conformation in the free protein.

**Figure 2.**
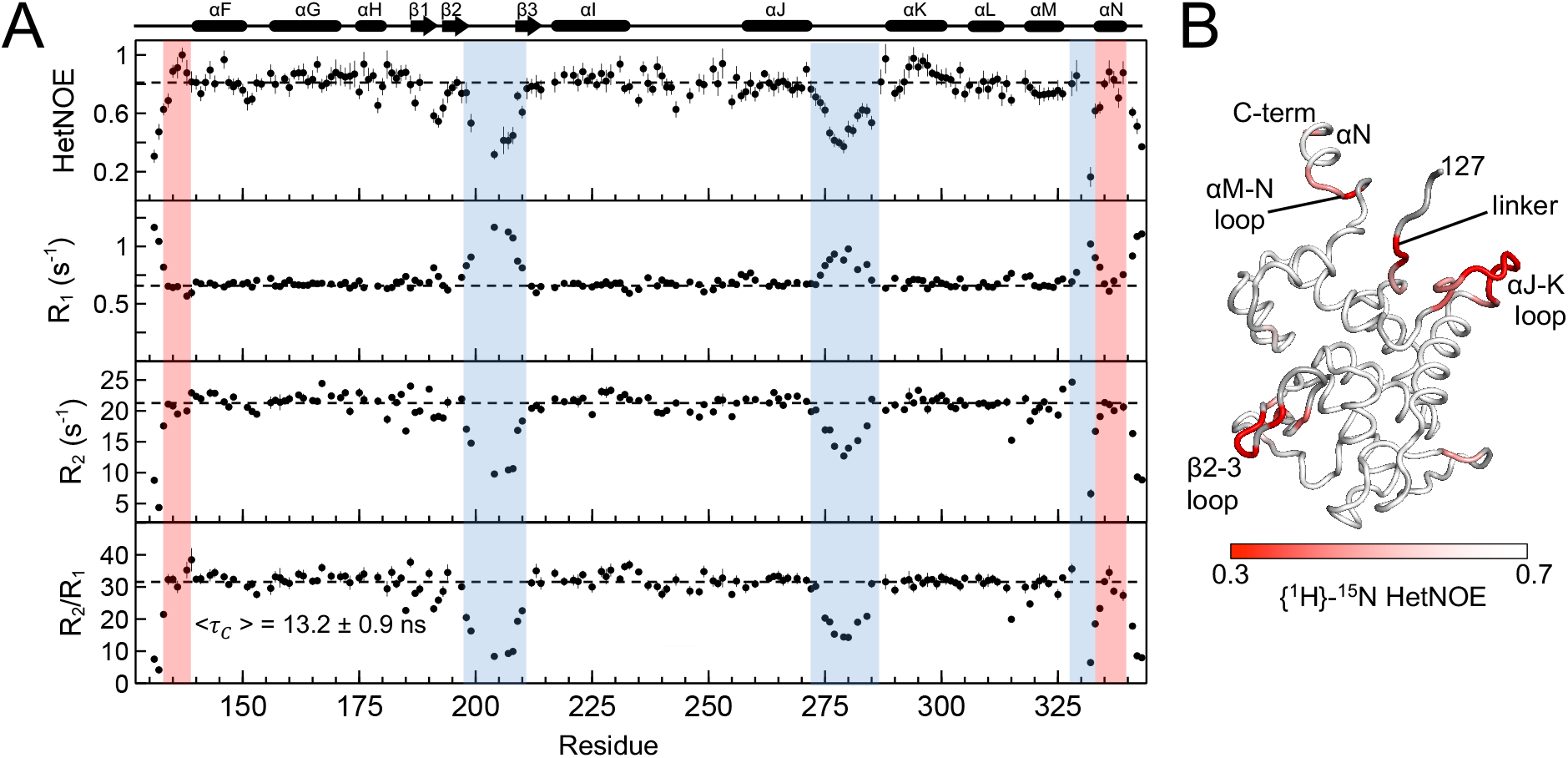
Cre^Cat^ loops β2-3, αJ-K and αM-N, but not helix αN, exhibit fast timescale dynamics. *(A)* {^1^H}-^15^N Heteronuclear NOE, ^15^N longitudinal R_1_ relaxation rate constant (s^−1^), ^15^N transverse R_2_ relaxation rate constant (s^−1^) and corresponding R_2_/R_1_ ratios for Cre^Cat^; 800 MHz; 298 K. Dashed lines in panels represent average values calculated excluding regions of fast timescale dynamics. The secondary structure elements of Cre^Cat^ synaptic complex crystal structure (PDB ID 2HOI, chain B) are shown above the plot for comparison. *(B)* {^1^H}-^15^N HetNOE values for Cre^Cat^ mapped onto the crystal structure (PDB ID 2HOI, chain B) as a linear gradient from red (= 0.3) to white (≥ 0.7). Residues with no data are shown in gray.

### Truncation of C-terminal helix αΝ results in widespread CSPs in the core of Cre^Cat^

In tetrameric DNA-bound crystal structures of Cre, the C-terminal helix αN of each protomer packs *in trans* in a cyclic (non-reciprocal) manner into a surface cavity on a neighboring protomer composed of residues from αF, αG, αH, β2-3 loop, αI, αK and αL regions (Fig. 1*A*). Since Cre is monomeric in the absence of DNA, the Van Duyne group had previously used crosslinking experiments to test the hypothesis that in free Cre the C-terminal region folds back to dock over its own *trans* cavity *in cis*; those experiments failed to detect intramolecular crosslinks^39^. Nevertheless, the ^15^N relaxation data (Fig. 2) would be consistent with a *cis* docking model in which rotational diffusion of αN is coincident with overall tumbling of the protein.

To test whether the C-terminal region of Cre^Cat^ might be docked *in cis*, instead of being extended for *trans* docking, we examined the effect of deleting the C-terminal residues on the NMR spectra. We constructed a C-terminal deletion lacking thirteen C-terminal residues including helix αN: Cre^Cat^ΔC (residues 127-330), obtained backbone resonance assignments from triple resonance NMR data, and compared the ^1^H-^15^N TROSY-HSQC spectra of WT Cre^Cat^ and Cre^Cat^ΔC (Fig. 3*A* and S4). Large chemical shift perturbations (CSPs) were observed for residues in the interdomain linker region (Q133-A136), helices αH, αK, αM and the αJ-K loop (Fig. 3*A*). Each of the regions in the protein core that show large CSPs is in excess of 20 Å from the center of helix αN in the synaptic structures. These findings are consistent with a *cis* docking arrangement of the C-terminus, and with the restricted internal motions inferred from ^15^N relaxation studies.

**Figure 3.**
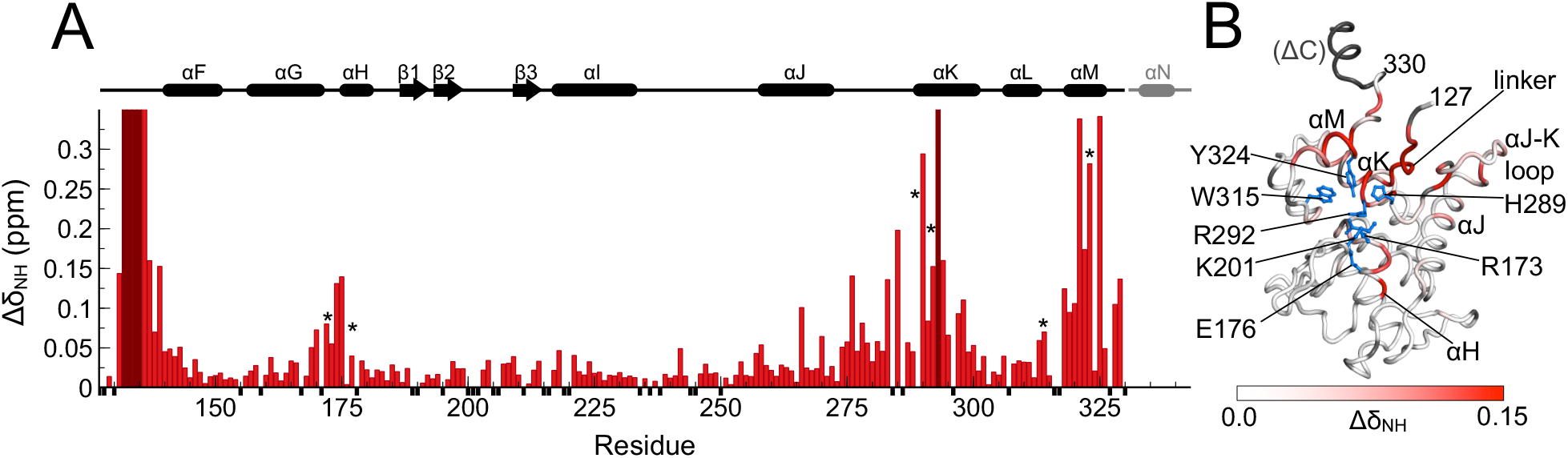
C-terminal deletion results in CSPs in the core of Cre^Cat^. *(A)* Per-residue amide CSPs [Δδ_NH_ = (Δδ_H_^2^ + Δδ_N_^2^/25)^1/2^] of Cre^Cat^ upon truncation of last 13 amino acid from the C terminus Δ(E331-D343) (red bars). The secondary structure elements of Cre synaptic complex crystal structure (PDB ID 2HOI, chain B) are shown above the plot. Asterisks indicate regions of high CSP containing active site residues R173, E176, H289, R292, W315 and Y324. Residues that showed largest CSPs or peak broadening beyond detection (assigned in Cre^Cat^ but unassigned in Cre^Cat^ΔC spectrum) are shown in maroon. Unassigned residues are indicated by small negative black bars. *(B)* Amide CSPs mapped onto the crystal structure (PDB ID 2HOI, chain B) as a linear gradient from white (= 0.0 ppm) to red (≥ 0.15 ppm). Sidechains of active site residues R173, E176, K201 (modeled using PyMOL), H289, R292, W315 and Y324 are shown in blue. Truncated C-terminal residues Δ(E331-D343) and unassigned residues are indicated in gray.

Remarkably, when the CSPs for Cre^Cat^ΔC are mapped to the crystal structure of the protein (Fig. 3*B*), the CSPs do not map to the synaptic *trans* docking surface of Cre. The region of the interdomain linker that shows large CSPs (Q133-A136) is adjacent to the DNA binding surface and was found to be largely rigid on the ps-ns timescale (Fig. 2). Helix αH contains active site residues R173 and E176, αJ-K loop and helix αK contain the active site residues H289 and R292, while helix αM contains the active site residue Y324.

These CSPs show that deletion of the C-terminal region alters the environment of residues surrounding the active site of Cre and part of its DNA binding surface. To ensure that the CSPs observed in these experiments are not an artifact of working with the catalytic domain Cre^Cat^, we compared spectra between full-length Cre and the same C-terminal deletion: CreΔC (residues 1-330). An overlay of ^1^H-^15^N TROSY-HSQC spectrum of full-length Cre on that of CreΔC showed the same CSPs for residues from the catalytic domain, and minimal perturbation of signals attributed to Cre^NTD^ (Fig. S5). Although CSPs can be due to either direct or indirect interactions^40,41^, the Cre^Cat^ΔC CSP data suggest that the C-terminal residues in Cre^Cat^ pack *in cis* over the active site and DNA binding surface of Cre^Cat^, not over the *trans* docking surface.

### PRE-NMR data support *cis* docking of C-terminal helix αΝ of Cre

To clarify whether the CSPs in the core of Cre^Cat^ domain were induced by direct interactions or via indirect allosteric effects, we attached a nitroxide spin probe at the extreme C-terminus of Cre^Cat^ and performed paramagnetic relaxation enhancement (PRE)-NMR experiments^42–44^. We constructed a variant of Cre^Cat^ (C155A/C240A/D343C) that enabled attachment of the spin probe to the C-terminal cysteine residue and used ^1^H-^15^N TROSY-HSQC spectra to verify that the mutations did not significantly perturb the structure of the protein (data not shown). The single cysteine mutant was then conjugated with a S-(1-oxyl-2,2,5,5-tetramethyl-2,5-dihydro-1H-pyrrol-3-yl) methyl methanesulfonothioate (MTSL) paramagnetic spin probe. Near 100% tagging efficiency was verified using MALDI mass spectrometry (data not shown). ^1^H-^15^N TROSY-HSQC spectra of the MTSL tagged-protein under oxidized (paramagnetic) and reduced (diamagnetic) conditions were obtained and a comparison of the peak intensities was used to determine per-residue PRE effects in free Cre^Cat^. In such a PRE-NMR experiment, the lone-pair electron on a paramagnetic spin probe is expected to induce distance-dependent line broadening for protons up to 25 Å away^45^.

A strong correspondence between the C-terminus-linked PRE effects and previously observed CSPs provide evidence for *cis* packing of the C-terminus over the active site and DNA binding surface of Cre. In addition to residues sequentially neighboring C343, we observed strong PRE effects from the C-terminal C343-MTSL spin probe to a part of the interdomain linker (E129-A136), helix αH (R173-A178), helix αJ (S257-I264), αJ-K loop (L284-R292) and helix αM (G314-Y324) (Fig. 4*A* and S6). To interpret the PRE effects in a structural context, we built a model of the MTSL tagged-protein using PyMOL^46^ with PDB ID 2HOI, chain B as the template, manually added the two missing C-terminal residues (G342, D343C), D343 substituted with cysteine, and attaching an MTSL moiety (coordinates from PDB ID 2XIU)^47^ to the C343 sidechain, and mapped the PRE data onto this Cre^Cat^ structural model (Fig. 4*C*: left). Each of the regions showing strong PRE effects in free Cre^Cat^ are at distances in excess of 25 Å as measured from the C343-MTSL-oxygen to backbone amide nitrogen atoms: further than ~ 42 Å, ~ 47 Å, ~ 51 Å, ~ 42 Å, and ~ 37 Å, respectively (Fig. S6 *A*).

**Figure 4.**
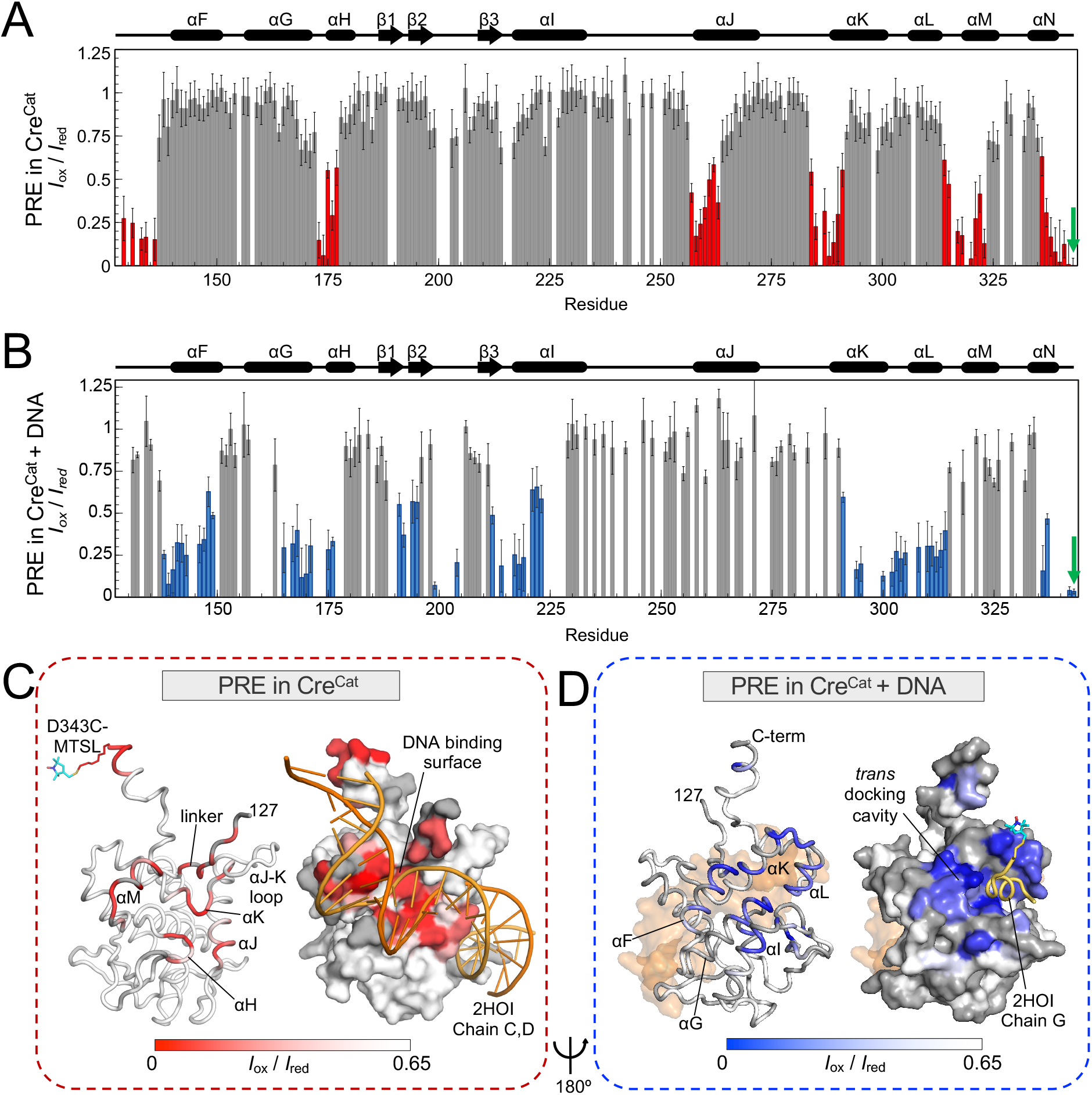
PRE-NMR data reveal a *cis* to *trans* docking conformational switch in the Cre^Cat^ C-terminus upon DNA binding. *(A)* Per-residue amide PRE-NMR normalized intensity ratios (*I*_ox_/*I*_red_) of free Cre^Cat^ C155A/C240A/D343C generated using an MTSL paramagnetic probe at C-terminal residue C343 (green arrow); strong PRE effects (*I*_ox_/*I*_red_ < 0.65) are shown in red. Secondary structure elements of Cre synaptic complex crystal structure (PDB ID 2HOI, chain B) are shown. Uncertainties are propagated from the signal-to-noise ratio of individual resonances. *(B)* Per-residue amide PRE-NMR normalized intensity ratios (*I*_ox_/*I*_red_) for Cre^Cat^ C155A/C240A/D343C bound to *loxP* half-site DNA generated using MTSL paramagnetic probe at C-terminal residue C343 (green arrow); strong PRE effects (*I*_ox_/*I*_red_ < 0.65) are shown in blue. Uncertainties are propagated from the signal-to-noise ratio of individual resonances. *(C) I*_ox_/*I*_red_ PRE values from free Cre^Cat^ mapped to the structure of a protomer in the crystal structure (PDB ID 2HOI, chain B; G342, C343-MTSL modeled) as a gradient from red (*I*_ox_/*I*_red_= 0) to white (*I*_ox_/*I*_red_= 0.65). Residues with no PRE data are shown in gray. Juxtaposed equivalent surface representation (same orientation) along with a portion of the *loxP* DNA, as seen in the crystal (PDB ID 2HOI, chains C and D) is shown in orange to illustrate the PRE effects mapping to the DNA binding surface. *(D) I*_ox_/*I*_red_ PRE values from the Cre^Cat^-DNA complex mapped to the structure of a protomer in the crystal structure (PDB ID 2HOI, chain B; G342, C343 modeled) as a gradient from blue (*I*_ox_/*I*_red_= 0) to white (*I*_ox_/*I*_red_= 0.65). Bound DNA (PDB ID 2HOI, chains C and D) is shown in pale orange. Residues with no PRE data are shown in gray. Juxtaposed equivalent surface representation (same orientation) along with C-terminal helix αN region from an adjacent MTSL-tagged protomer (PDB ID 2HOI, chain G: E331-D341 and G342, C343-MTSL modeled) is shown in yellow to illustrate the PRE effects mapping to protein-protein *trans* docking surface cavity. Orientation of Cre^Cat^ is rotated approximately 180° between Fig. 4*C* and *D*.

To rule out the possibility that the observed PRE effects were due to intermolecular interactions, we performed an intermolecular PRE experiment by recording ^1^H-^15^N TROSY-HSQC spectra of a 1:1 mixture of [U-^15^N]-Cre^Cat^ (non-tagged) with MTSL tagged-unlabeled Cre^Cat^ C155A/C240A/D343C, under paramagnetic and diamagnetic conditions. Because the ^15^N labels and MTSL tags are on different molecules any PRE effects must arise from intermolecular interactions (Fig. S7 *A*)^48^. Only a few sparse residues located in the *trans* docking surface of Cre showed intermolecular PRE effects, suggestive of very weak transient intermolecular interaction (Fig. S7 *B* and *C*). Thus, the strong intramolecular PRE effects in the 1H-^15^N TROSY-HSQC amide spectrum of free Cre^Cat^ strongly support a model in which the C-terminal region makes a *cis* interaction with its own DNA binding surface (Fig. 4*C*: left and right).

### PRE-restraints yield a *cis* docked model of the C-terminus of Cre, consistent with CSPs

To determine whether the covalent structure of Cre^Cat^ would allow *cis* docking of helix αN region without distorting other structural elements, we performed structure refinement using PRE-derived distances as restraints. Analysis of the free Cre^Cat^ PRE *I_ox_*/*I*_*red*_ values and ^15^N R_2_ relaxation rates yielded 124 restraints (Table S1). To account for a distribution of spin probe positions, lower bounds were set to 5 Å below calculated distances while the upper bounds were set to 10 Å higher than the calculated distances. ROSETTA energy minimization was performed using upper and lower distance bounds, assuming a flexible C-terminus, β2-3 loop and αJ-K loop regions; crystallographically observed secondary structure elements were enforced. Members of the PRE-derived Cre^Cat^ ensemble (Fig. 5*A*) adopt stereochemically robust *cis* docked C-terminal conformations that differ significantly from the conformation observed in synaptic complexes (Fig. 5*B*). These structures are consistent with CSP data, even though those were not used during refinement (Fig. 5*C*). Difference Cα-Cα contact map between the top ten conformers and the synaptic crystal structure (PDB ID 2HOI, chain A) show that the average changes in C-terminal inter-residue distances when *cis* docked, ranged in values as large as 25 Å (Fig. 5*D*).

**Figure 5.**
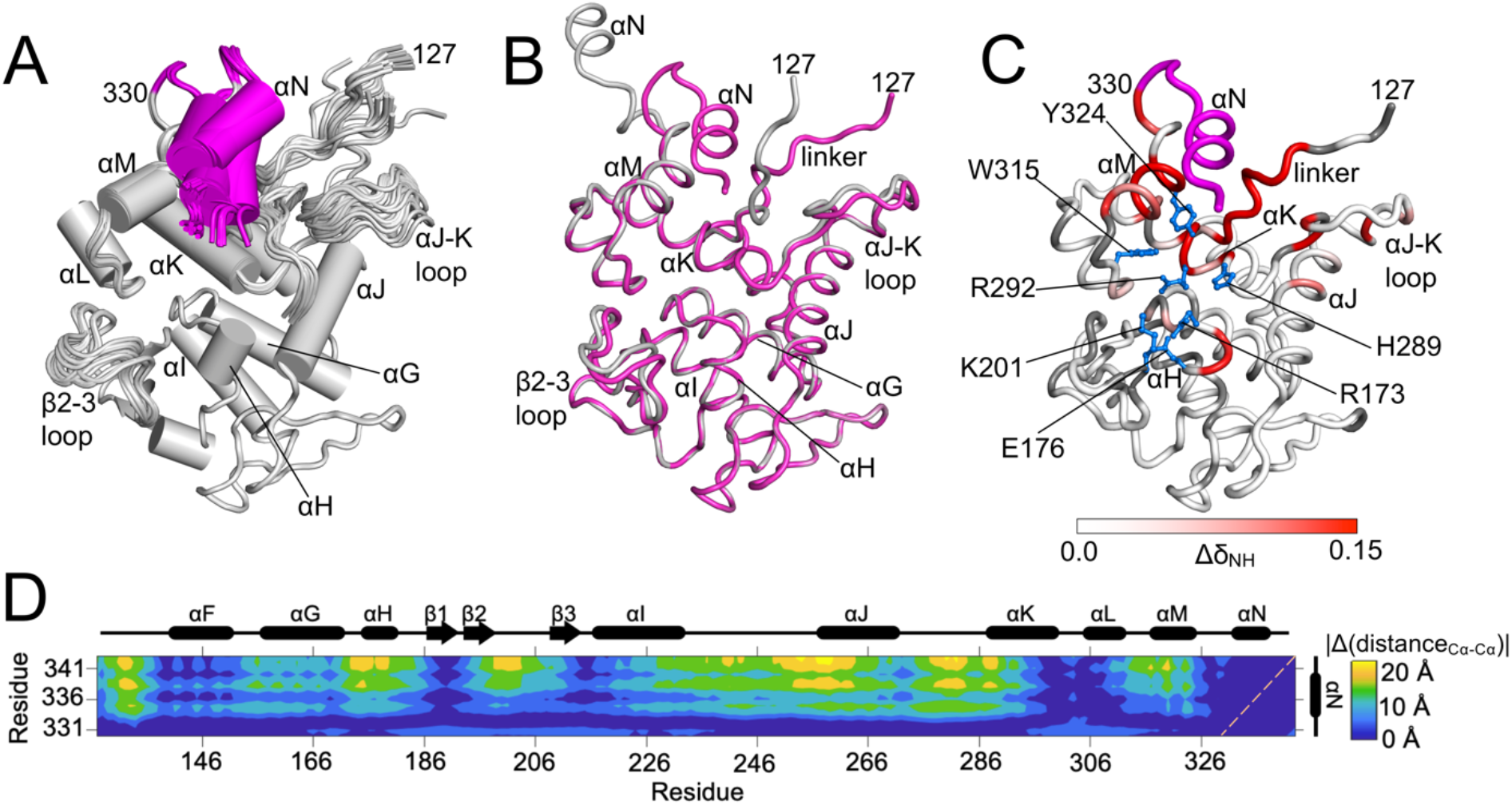
PRE-derived ensemble models of free Cre^Cat^ shows *cis* docking of C-terminus over the Cre^Cat^ active site. *(A)* Top fifty PRE-derived ROSETTA models of free Cre^Cat^ showing *cis* docking of C-terminal region (E331-D343; in magenta). *(B)* Overlay of lowest ROSETTA energy member of the PRE-derived ensemble model (magenta) and the crystal structure protomer (PDB ID 2HOI, chain A) (gray). *(C)* CSPs in free Cre^Cat^ upon C-terminal truncation Δ(E331-D343) (magenta) mapped onto the lowest energy member of the PRE-derived ensemble model as a linear gradient from white (0.0 ppm) to red (≥ 0.15 ppm). Unassigned residues are shown in gray (as in Fig. 3*B*). *(D)* Difference in the inter-residue Cα-Cα contact map between the top ten PRE-derived models and the DNA bound tetrameric crystal structure (PDB ID 2HOI, chain A) (absolute values of changes in inter-residue distances |Δdistance_Cα-Cα_| (Å) are shown by a three-color heatmap from blue to yellow). The secondary structure elements of Cre synaptic complex crystal structure (PDB ID 2HOI, chain B) are shown along the axes. Diagonal is shown as dashed line (orange).

### DNA binding perturbs the C-terminal region in Cre^Cat^

Coincidence of the DNA binding site and *cis* docking surface of the Cre^Cat^ C-terminal region led us to examine the effects of binding to a *loxP* DNA half-site substrate (Fig. 1*C*), on the protein C-terminus. The ^1^H-^15^N TROSY-HSQC spectrum of the Cre^Cat^-DNA half-site complex showed CSPs in residues corresponding to the Cre-DNA interaction surface, comprising of structural elements involved in DNA binding and the active site regions. Moreover, large CSPs (and broadening of resonances) were also observed in the C-terminal residues E331-D343, including helix αN (Fig. 6*A* and *B*). The CSPs mapped on the crystal structure (Fig. 6*C*) show that *loxP* DNA half-site binds to Cre^Cat^ in the manner predicted from available tetrameric structure models, but DNA binding also induces a change in the environment of the C-terminal residues, resulting in the observed CSPs.

**Figure 6.**
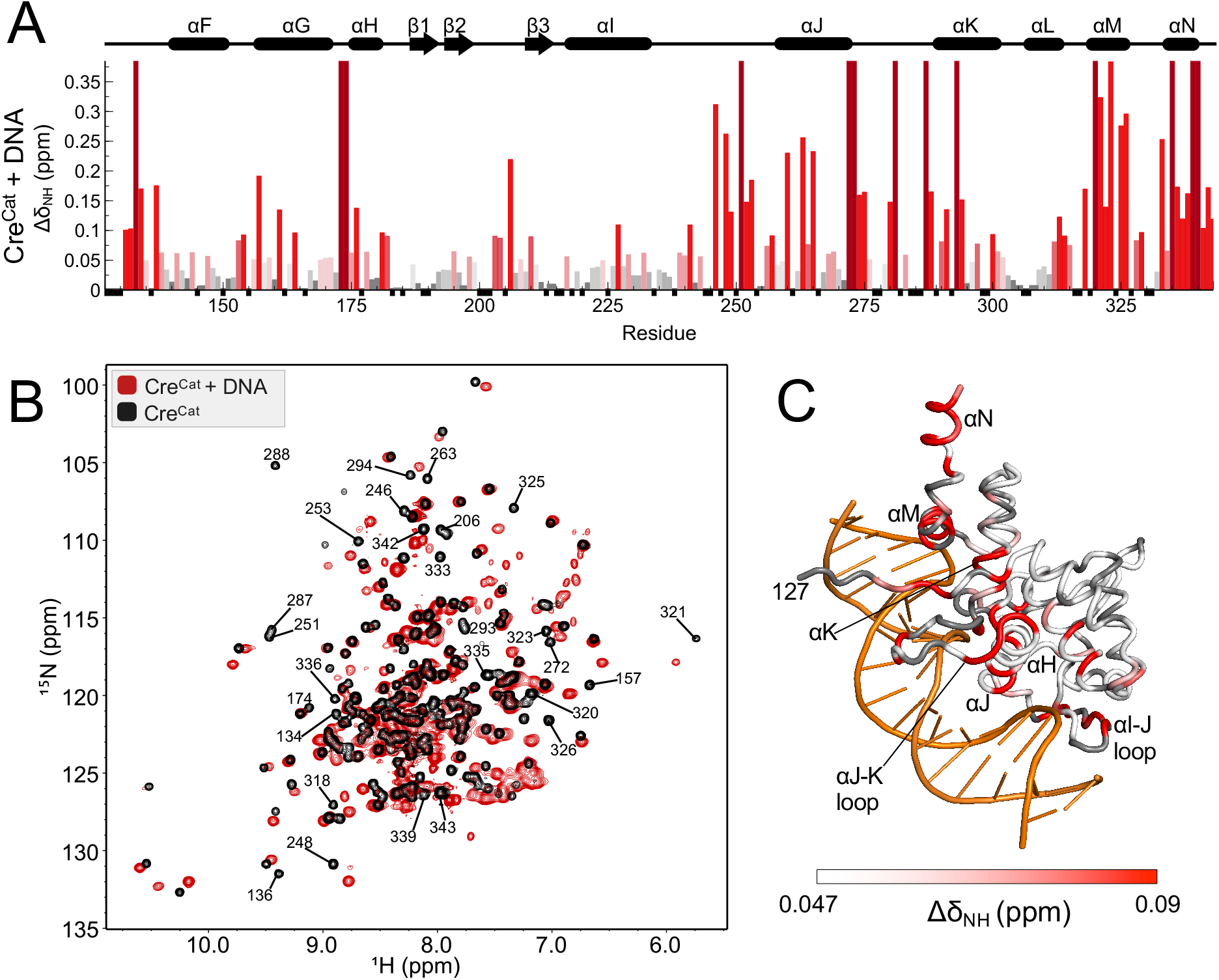
C-terminal helix αN region of Cre^Cat^ is perturbed upon binding to *loxP* DNA *(A)* Per-residue amide CSPs [Δδ_NH_ = (Δδ_H_^2^ + Δδ_N_^2^/25)^1/2^] of Cre^Cat^ upon binding to *loxP* half-site hairpin DNA with a linear color ramp from gray (= 0 ppm) to red (≥ twice the Δδ SD (0.047 ppm)). Largest CSPs, or those that resulted in peak-broadening beyond detectability (assigned in free Cre^Cat^ but unassigned in the Cre^Cat^-DNA complex) are shown in maroon. Unassigned residues are indicated by small negative black bars. Secondary structure elements of Cre synaptic complex crystal structure (PDB ID 2HOI, chain B) are shown above the plot. *(B)* Overlay of ^1^H-^15^N TROSY-HSQC spectra of free Cre^Cat^ (black) and bound to a DNA half-site (red) with select residue assignments indicated (> 3 SD ppm). *(C)* CSPs mapped onto the crystal structure (PDB ID 2HOI, chain B) as a linear gradient from white (≤ 1 SD ppm) to red (≥ 2 SD ppm). Unassigned residues are shown in gray.

### PRE-NMR data reveal a DNA binding-induced conformational change in the Cre C-terminus

Like for the free protein, we used PRE-NMR to determine the conformational space populated by the C-terminus of Cre upon binding to *loxP* DNA half-site. PRE effects were measured using MTSL tagged-Cre^Cat^ C155A/C240A/D343C bound to the *loxP* half-site hairpin DNA. PRE effects were remarkably different in the DNA complexes in comparison to free Cre^Cat^ (Fig. 4*B* and S8). Low *I_ox_*/*I*_*red*_ ratios were observed for helix αF (E138-M149), helix αG (G165-E176), β2 and β2-3 loop (G191-I195, G198 and V204), β3-αI region (A212-R223), and helices αK-αL region (A291-G314). These regions together form the *trans* docking cavity that accommodates the C-terminus of neighboring Cre protomers in synaptic complexes (Fig. 1*A* and 4*D*: left and right). This change in PRE pattern (Fig. 4*A* versus 4*B*) clearly identifies a DNA-induced conformational change that involves displacing the C-terminus from the DNA binding surface, to enable new protein-protein interactions; however, they do not establish whether those new interactions occur *in cis* (intramolecular), or *in trans* (intermolecular).

Intermolecular PRE-NMR studies with mixed isotopic labeling were then used to clarify whether the PRE effects observed in the presence of DNA arise from intra or intermolecular interactions. We mixed DNA-bound non-tagged [U-^15^N]-Cre^Cat^ and MTSL-tagged unlabeled Cre^Cat^ C155A/C240A/D343C at a 1:1 molar ratio, to study possible intermolecular interactions in Cre^Cat^ bound to DNA. Because these mixed samples have the potential to interact in four different relative orientations (Fig. S9 *A*), only one of which would generate measurable PRE effects (i.e., MTSL tagged-unlabeled Cre^Cat^ C155A/C240A/D343C with C-terminus docking onto a [U-^15^N]-Cre^Cat^), the experiment would be predicted to yield weaker PRE effects in comparison to the experiments with uniformly labeled and MTSL tagged-Cre^Cat^-DNA complex (Fig. 4*B*); although observation of such effects in the same regions of the protein would support *trans* interactions. Despite the resonances being broader and data quality being generally lower, we indeed observed strong PRE effects consistent with docking of DNA-bound Cre C-terminal region over the *trans* docking surface (Fig. S9 *B* and *C*); these effects *must* arise from intermolecular interactions because of the labeling pattern used. Yet, a comparison to the similarly strong PRE effects in uniformly tagged and labelled Cre^Cat^-DNA complex studies (Fig. S9 *C* versus Fig. 4*B*) illustrates the possibility of a population of DNA-bound Cre C-terminus docking *in cis* over its own *trans* docking cavity, while another population extends out to dock *in trans* into the same cavity on another DNA-bound Cre molecule.

## DISCUSSION

Cre recombinase has emerged as an important tool in molecular and cellular biology and has several features that make it an attractive reagent for gene editing^7,49,50^. Reaching that potential requires thorough characterization of its mechanism for DNA site selection, and control of its DNA cleavage and recombination activity. High-resolution structural studies of Cre, largely limited to crystal structures of DNA bound tetrameric synaptic complexes, have suggested a role for conformational plasticity in facilitating the steps within the reaction mechanism. We set out to characterize the solution behavior of Cre in its free and DNA bound states in order to understand the link between protein dynamics and regulation of Cre assembly and activity. While solution NMR studies yielded some results consistent with previous understanding of this prototypical member of the tyrosine recombinase family, other surprising results call for a revised view of the reaction mechanism.

Solution NMR data showed Cre^Cat^ to be independent of Cre^NTD^. The well-dispersed signals in the ^1^H-^15^N TROSY-HSQC NMR spectrum of Cre^Cat^ are nearly superimposable on that of full-length Cre, except for the signals attributable to N-terminal domain Cre^NTD^ (Fig. S1 *B*). The domains’ structural independence could imply variability in inter-domain orientations that would be expected to prevent formation of suitable crystal lattices for x-ray diffraction. It however facilitated detailed NMR studies with the Cre^Cat^ domain, wherein reside the active site residues and sequence specificity determinants, focused on testing the role of protein dynamics in regulation of Cre function^33–36^.

Our findings of the protein dynamics in specific regions within Cre^Cat^ revealed unique spectral signatures linked to their functional roles. Underlying flexibility in Cre^Cat^ studied by nuclear spin relaxation measurements (^15^N R_1_, R_2_ and {^1^H}-^15^N HetNOE), were consistent with the monomeric 217-residue protein construct (τ_*C*_, 13.2 ns). A portion of the linker, the β2-3 loop, the αJ-K loop and the αM-N loop each exhibit higher R_1_, lower R_2_, and reduced HetNOE values, indicative of fast internal motions. Within the Cre_4_-*loxP*_*2*_ tetrameric synaptic complexes, conformational differences in the β2-3 loop and the C-terminal regions (including helix αN) that form structural bridges that link adjacent protomers, and in helix αM bearing the catalytic tyrosine, distinguish the catalytically “active” and “inactive” Cre protomers^5,16,22^. Flexibility of the β2-3 loop observed in free Cre^Cat^ could be expected to prevent stabilization of the active site geometry in uncoordinated Cre protomers. The alternating conformations of the loop in tetrameric synaptic complexes indicates that this flexibility will be still retained, thereby enabling its regulatory role over adjacent active site structures. The protein dynamics observed in the αJ-K loop in the free protein may serve to facilitate DNA binding and could thereafter be predicted to undergo quenching since this region does not show structural plasticity between the various Cre reaction intermediates. The surprise within the protein dynamics analysis lies in the observation that while residues flanking C-terminal helix αN exhibit flexibility on the ps-ns timescale, αN itself does not, and instead exhibits relaxation rates consistent with overall tumbling of the protein, counter to the expectation that the Cre C-terminus would flexibly adopt a range of extended conformations to facilitate capturing an adjacent protomer via *trans* docking contacts.

An unexpected C-terminal *cis* docking interaction in the proximity of the free Cre active site was revealed from our highly correlated CSP and PRE-NMR experiments. Given the known *trans* docking site, and prior incongruities regarding *cis* or *trans* cleavage by Cre and other recombinases^51^, a logical presumption was that in free Cre, helix αN might dock *in cis* in the same site it occupies when assembled *in trans* in synaptic complexes; indeed, such a premise was previously tested experimentally using disulfide crosslinking strategies^39^. Were that the case, one would predict that deletion of helix αN would result in CSPs for residues flanking that *trans* docking surface cavity. Instead, we observed the largest CSPs for residues on the other side of the protein – namely, the DNA binding and active site surfaces (Fig. 3). Since CSPs can arise from both proximity and induced conformational changes, we employed distance-dependent effects in PRE-NMR experiments to conclusively demonstrate the proximity of the protein C-terminus to the active site and DNA binding regions (Fig. 4*A*). Consistent with size exclusion chromatography (Fig. S1 *A*), NMR relaxation (Fig. 2) and previous sedimentation velocity experiments^17^ that show unbound Cre to be monomeric in solution, the absence of strong intermolecular PREs (Fig. S7) indicates that docking of the C-terminus of free Cre onto the DNA binding surface indeed occurs *intra-*molecularly, *in cis*. *Cis*-docked models generated using distance bounds derived from the PRE data satisfy the restraints, covalent structure, and are in excellent agreement with the CSPs obtained upon deletion of the C-terminal region (Fig. 5). A comparison between the free Cre^Cat^ ensemble model and the Cre^Cat^ structure from crystallographic DNA bound synaptic complex protomers shows changes in inter-residue contacts as large as 20-25 Å (Fig. 5*D*). Thus, these data provide strong evidence that in the absence of DNA the C-terminal residues dock *in cis* over the DNA binding and active site surfaces, in an apparent auto-inhibitory state.

A Cre structure in which the C terminus is docked *in cis* over the active site would seem incompatible with a DNA-bound state – so, how is this conundrum resolved? PRE-NMR data on the Cre^Cat^-DNA complex show that the C-terminus of Cre^Cat^ is displaced from the *cis* docking surface when DNA binds there, since PRE effects are instead observed at the *trans* docking cavity (Fig. 4*B*). This *loxP* binding-induced pattern of PREs at the *trans* docking site clearly indicates relocation of C-terminal helix αN, but does not clarify whether docking occurs *in cis*, or *in trans* to a nearby Cre molecule. The use of a DNA *loxP* half-site substrate would be expected to prevent assembly of pre-synaptic complexes with two Cre molecules bound to DNA; however, PRE effects can be observed even when interactions are transient^52^. Although partially attenuated, intermolecular PRE effects were indeed observed at the same *trans* docking cavity in experiments where the NMR signals from DNA-bound Cre^Cat^ arise from protein molecules not tagged with MTSL (Fig. S9). This attenuation could result from the fact that only one-fourth of protein-protein contacts within DNA-bound complexes can produce *intermolecular* PRE effects (i.e., when C-terminus from MTSL-tagged un-labeled Cre^Cat^ docks *in trans* over non-tagged [U-^15^N]-Cre^Cat^ when bound to DNA; Fig. S9 *A*), or may indicate that in the presence of DNA, *intramolecular cis* docking over the protomer’s own *trans* docking cavity is also possible. However, molecular modeling calculations starting from the DNA-bound crystal structure (data not shown) indicated that the C-terminal helix αN region could not pack over its own *trans* docking cavity without significant conformational strain, involving additional remodeling that would be inconsistent with the NMR CSP data (Fig. 5). We conclude that in the absence of DNA, the C-terminal residues spanning helix αN occupy the DNA binding surface of Cre, and that binding to DNA produces a large conformational change that enables productive *trans* protein-protein interactions with adjacent Cre molecules.

The rigidity and apparent auto-inhibitory conformation observed in C-terminal residues in free Cre could be expected to disfavor oligomerization of the protein prior to *loxP* DNA binding and play a role in selectively inhibiting spurious uncoordinated DNA cleavage and recombination activity. Although it is unclear whether the *cis* docking conformation of C-terminal region in free Cre serves additional roles, for example by obstructing inadvertent binding to non-cognate DNA substrates, our data suggests that the conformational rearrangements following the displacement of *cis* docking C-terminal upon *loxP* DNA half-site binding, comprise the mechanistic step that could trigger Cre protomer stepwise assembly on *loxP* through protein-protein interactions via the now exposed C-terminal residues.

These studies expand our understanding of the regulatory role played by C-terminal residues in Cre^5,16,17^. DNA-binding associated conformational changes mediated by C-terminal regions of Cre and related recombinases have been implicated in catalytic activation and initiation of oligomerization^2,5,28^. However, the structural basis for Cre to undergo stepwise assembly on the DNA recognition sites have remained poorly understood due to lack of structural information on pre-synaptic reaction intermediates. The features of Cre and Cre-*loxP* complexes presented here provide insight into both the unliganded state of Cre, as well as the mechanism of initiation of protein-protein interactions that lead to stepwise assembly of recombinogenic intasomes. Knowledge gaps remain, including understanding of precise structural rearrangements that accompany the *cis* to *trans* switch in pre-synaptic Cre recombinase. Nevertheless, these investigations provide a new NMR perspective on Cre recombinase and represent a transformative step towards understanding the structure, function and mechanism of this important and widely used gene editing tool.

## MATERIALS AND METHODS

### Protein expression, purification, and site-directed mutagenesis

The deletion construct containing the catalytic domain and the interdomain linker (Cre^Cat^, residues 127-343) was prepared using QuikChange Site-Directed Mutagenesis Kit (Agilent) from a pET21A vector (Novagen) encoding WT Cre Recombinase (provided by Dr. Gregory Van Duyne, U. Penn). Further truncation and point mutants of Cre^Cat^ construct were prepared using the Q5 Site-Directed Mutagenesis Kit (NEB). Protein expression from *E. coli* BL21(DE3) cells was carried out by growing freshly transformed cells (using Ampicillin antibiotic, 100 mg/L) in LB media (natural abundance Cre) or M9 minimal medium (isotopically labeled [U-^15^N] and [U-13C,^15^N] Cre) supplemented with 1 g/L [^15^N]-ammonium chloride (Cambridge Isotopes) as the sole nitrogen source or 2 g/L [^13^C]-glucose (Cambridge Isotopes) as the sole carbon source in addition to [^15^N]-ammonium chloride. The cells were grown (shaking at 220 rpm, 37 °C) to an OD_600_ of 0.6-0.8 and induced with 0.75 mM isopropyl β-D-1-thiogalactopyranoside (Gold Biotechnology) for 12 h. After harvesting the cells (4000 g, 15 min, 4 °C) and sonication, the Cre constructs in 40 mM tris, 100 mM NaCl, pH 7.0 buffer containing 1 protease inhibitor cocktail tablet (Roche), were purified using 5 ml SPFF cation exchange column (GE Healthcare) with a 100 ml gradient of 0.1 to 1 M NaCl. The protein fractions (eluted at ~ 0.4 M NaCl) were combined and diluted four times with low salt buffer 40 mM tris, 100 mM NaCl pH 7.0 and further purified on a 5 ml Heparin affinity column (GE Healthcare) with a 100 ml gradient of 0.1 to 1.5 M NaCl. Protein fractions (eluted at ~ 1 M NaCl) were then purified using a HiLoad 16/60 Superdex 75 (GE Healthcare) column (SEC) in 40 mM tris, 500 mM NaCl pH 7.0 buffer. Purified protein concentrations were determined using UV-Vis spectroscopy (ε for Cre^Cat^ = 23,950 M^−1^cm^−1^ at 280 nm). Typical yields obtained were ~20-25 mgs protein per L of culture.

### Cre^Cat^-DNA complex NMR sample preparation

*loxP* half-site hairpin was formed from single-stranded DNA oligos (IDT) containing recombinase binding elements along with additional GC bps at ends, symmetrized spacer bps and a GAA hairpin^53^: 5’-GCATAACTTCGTATAGCATATGCGAAGCATATGCTATACGAAGTTATGC-3’ The lyophilized DNA oligo was resuspended in H_2_O and heated at 95 °C for 15 mins and immediately cooling on ice. To avoid precipitation, Cre^Cat^ and *loxP* DNA hairpin at low concentrations (< 50 μM) in high salt buffer (10 mM Tris, > 500 mM NaCl, pH 7.0) were mixed by stepwise titration of the protein into DNA. The solution was then dialyzed into low salt NMR buffer (10 mM tris, 15 mM NaCl, 0.02% NaN_3_, pH 7.0) at 4 °C. The protein-DNA complex sample was then concentrated using 500 μL, 1 kDa centrifugal filters (VWR).

### NMR data collection

Purified [U-^15^N]- or [U-^15^N,^13^C]-Cre^Cat^ samples were concentrated to 0.5 - 1.0 mM after dialysis into buffer containing 10 mM Tris, 100 mM NaCl, 0.02 % NaN_3_, pH 7.0. When required, the samples were exchanged into 10 mM d_11_-Tris (Sigma), 100 mM NaCl, 0.02 % NaN_3_, pH 7.0 buffer, using Sephadex PD10 columns (GE Healthcare). 5-10 % (v/v) D_2_O (99 %) and 0.66 mM DSS were added to the NMR samples. NMR data were recorded on Bruker Avance III HD spectrometers operating at 850, 800 or 600 MHz equipped with a 5 mm triple resonance cryoprobes and z axis gradients, at 25 °C. Data were processed with NMRPipe^54^ or NMRFx^55^ and visualized using NMRView^56^. Typical ^1^H-^15^N TROSY-HSQC 2D correlation spectra were recorded using 16 - 24 scans and 2048 × 128 data points.

### NMR chemical shift assignment and perturbation mapping

For backbone chemical shift assignments, TROSY triple-resonance spectra HNCO, HNCA, HN(CO)CA, HN(CA)CO, and non-TROSY CBCA(CO)NH and HA(CO)NH were recorded on purified [U-^13^C,^15^N] Cre^Cat^ sample. Backbone assignment was achieved using NMRView aided by PINE^57^ and CARA^58^. The assignments have been deposited in the Biological Magnetic Resonance Data Bank, www.bmrb.wisc.edu (accession no. 50270).

CSPs were calculated from differences in peak positions using:

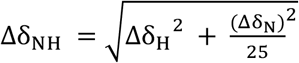

where Δδ_H_ is the chemical shift difference in the ^1^H dimension and Δδ_N_ is the chemical shift change in the ^15^N dimension. Corrected standard deviations were determined by calculating the SD of CSPs of all assigned residues, removing CSP data with > 3 SD and recalculating the SD.

### ^15^N relaxation measurements

Backbone amide relaxation measurements were performed at 800 MHz proton Larmor frequency using [U-^15^N] Cre^Cat^ at a concentration of 800 μM in 10 mM Tris, 100 mM NaCl, 0.02% NaN_3_, pH 7.0 buffer (5 % D_2_0 v/v). TROSY versions of R_1_ and R_2_ experiment data were acquired using recycle delay of 2 seconds between experiments, and the following relaxation delays for R_1_: 0, 560, 1120 and 1680 ms with repeats and R_2_: 2, 24.2, 48.2, and 72.2 ms. The R_1_ and R_2_ values were obtained by fitting the intensity of peaks to exponential decay function and errors were determined by relaxation curve fitting. 80-90 % of the amides were analyzed, after excluding those with significant resonance overlap. {^1^H}-^15^N HetNOE values were obtained by recording spectra with and without a ^1^H pre-saturation period (8 s), in which ^1^H signals were saturated using a train of 90° pulses, applied before the start of experiment. Non-overlapping assigned peaks were analyzed and the intensity ratio were determined using the HetNOE analysis tool within NMRViewJ; uncertainties were obtained from standard deviation of noise in the spectra.

### Paramagnetic Relaxation Enhancement (PRE)-NMR

We constructed a single-cysteine Cre^Cat^ variant C155A/C240A/D343C to avoid unintended paramagnetic tagging of the native cysteines and thereby enable probing just the C-terminus. S-(1-oxyl-2,2,5,5-tetramethyl-2,5-dihydro-1H-pyrrol-3-yl) methyl methanesulfonothioate (MTSL) tagging of Cre^Cat^ C155A/C240A/D343C was achieved by following a published protocol^59^. Briefly, a 200 mM MTSL (Toronto Research Chemicals) stock was made by adding 189 μL of acetonitrile to 10 mg of MTSL (stored at −20 °C, protected from light). DTT was added to purified protein in high salt conditions (10 mM Tris, 500 mM NaCl, 5 mM DTT, pH 7.0) to reduce possible disulfide bonds. The reducing agent was then removed by rapid buffer exchange into 10 mM Tris, 500 mM NaCl, pH 7.0 buffer using a PD10 desalting column (GE Healthcare). The sample was collected into exchange buffer containing ten-fold molar excess of MTSL and stirred at room temperature overnight. Excess MTSL was removed after reaction by buffer exchange with a second PD10 column, followed by dialysis (MWCO 3 kDa) into NMR buffer. Near 100 % MTSL tagging of the protein was verified using MALDI MS measurements as indicated by an increase in mass by 186 Da (weight of attached probe). Final sample concentrations ranged between 100-200 μM at a volume of ~ 500 μL.

For PRE-NMR measurements, ^1^H-^15^N TROSY-HSQC spectra were recorded on the same MTSL tagged-Cre^Cat^ C155A/C240A/D343C sample before (paramagnetic) and after (diamagnetic) reduction of the spin probe by treatment with five-fold molar excess of sodium ascorbate (250 mM stock used for minimal sample dilution) for three hours at room temperature. Peak intensities were measured and normalized using intensities of residues (D153, E222, V230) > 45 Å away from C-terminus in PDB ID 2HOI, chain B. For MTSL probes, *I_ox_*/*I*_*red*_ ratios between 0 and 1 indicate that the distance of probe to proton is within 13-25 Å. Uncertainties in *I_ox_*/*I*_*red*_ values were propagated from signal-to-noise ratio of each resonance in the two spectra using:

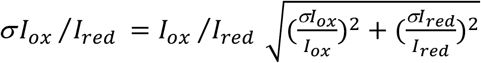

where σ*I_ox_*/*I*_*red*_ is the calculated error in *I_ox_*/*I*_*red*_ ratio, σI_ox_ is the standard deviation of the noise in the MTSL oxidized spectrum, and σI_red_ is the standard deviation in the reduced spectrum.

For the intermolecular PRE studies, [U-^15^N]-Cre^Cat^ (non-MTSL-tagged) was mixed with MTSL tagged-natural abundance Cre^Cat^ C155A/C240A/D343C, at a 1:1 molar ratio in NMR buffer (10 mM Tris, 100 mM NaCl, pH 7.0) at 25 °C. For similar studies with the protein-DNA complex, the complex samples with/without MTSL tags were prepared separately and mixed at a molar ratio of 1:1.

### PRE-restrained modeling using ROSETTA

PRE derived-distance constraints for use in structural modeling were generated using the approach of Battiste and Wagner^42^. Briefly, per residue R_2_^sp^ values (paramagnetic relaxation rate due to the spin tag) were obtained from:

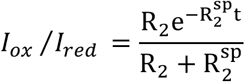

where R_2_ is the transverse relaxation rate of each amide and t is the acquisition time in the proton dimension. R_2_^sp^ values were then used to obtain the approximate distance, r (Å) between the spin tag to the amide proton of each residue:

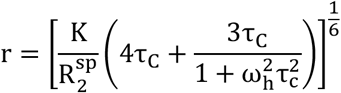

where τ_*C*_ (s) is per-residue correlation time of amide protons determined from R_2_/R_1_ relaxation ratios, ω_h_ (Hz) is the Larmor frequency of each amide proton, and K is a constant (1.23 × 10^−32^ cm^6^ sec^−2^) that encapsulates the spin properties of the MTSL tag^42^.

Distance restraints were used in a bounded form (typical for NOE constraints). Due to the relative imprecision of this method of constraint distance calculation^44,60^, residues with an *I_ox_*/*I*_*red*_ value less than 0.8 were assigned a lower bound of calculated *r -* 5 Å and an upper bound was set to *r* + 10 Å. Residues with an *I_ox_*/*I*_*red*_ value greater than 0.8 were assigned a lower bound of 20 Å and no upper bound. Constraints for 124 residues in Cre^Cat^ were obtained wherein data (*I_ox_*, *I_red_*, R_2_, ω_*h*_, and τ_*C*_) for each amide proton was available.

ROSETTA energy minimization was performed using the relax application^61–65^. Chain A of PDB ID 2HOI (residues 127-341, and G342 and D343C modeled) was used as the input structure with the D343C mutation. A move map file retained secondary structure of Cre^Cat^ along with the following impositions: the β2-3 and αJ-K loops and interdomain linker residues (A127-A134) were allowed to have backbone φ, ψ, and sidechain χ_1_ torsional freedom based on flexibility observed via ^15^N R_2_/R_1_ data. C-terminal E331-C343 were also allowed to sample backbone and χ_1_ angles allowing movement of helix αN through space. Other residues were only allowed to undergo sidechain minimization. Constraints were used to generate 100 initial structures subsequently scored using the ref2015_cst weights function^66^. The ten structures with the lowest *atom_pair_constraint* energy were selected as the input structures for a second round of minimization wherein ten structures were generated for each input structure. The resulting 100 structures constitute the PRE-NMR constrained ensemble model of free Cre^Cat^.

Coarse grained contact maps were generated using MATLAB by calculating backbone inter-residue Cα-Cα distances from PDB files (2HOI, chain A residues 127-341 and G342 and D343C modeled, and top ten lowest energy members of free Cre^Cat^ PRE-derived ensemble). Absolute values of change in inter-residue distances between the contact maps were determined by subtracting one distance matrix from the other.

## AUTHOR CONTRIBUTIONS

Author contributions: A.U., C.A. and D.K.Y. and M.P.F. designed experiments; A.U., C.A., D.K.Y and D.P. performed research; A.U., D.K.Y., C.A., K.S. and M.P.F. analyzed and interpreted data; and A.U. and M.P.F. wrote the manuscript.

## ACKNOWLEDGEMENTS

We thank Dr. Chunhua Yuan, Dr. Alex Hansen and Dr. Arpad Somogyi of the Ohio State University Campus Chemical Instrumentation Center for assistance with NMR experimental setup and mass spectrometry analysis and Dr. Eric Danhart for guidance with NMR data processing. We thank Dr. Gregory Van Duyne (U. Penn) for providing the plasmids encoding Cre. This study was supported by NIH grant R01 GM122432 (to M.P.F), NSF MCB grant 0092962 (to M.P.F), and NIH/NIGMS grant 1P41GM111135 (to NMRBox).

## SUPPLEMENTARY INFORMATION

**Figure S1.**
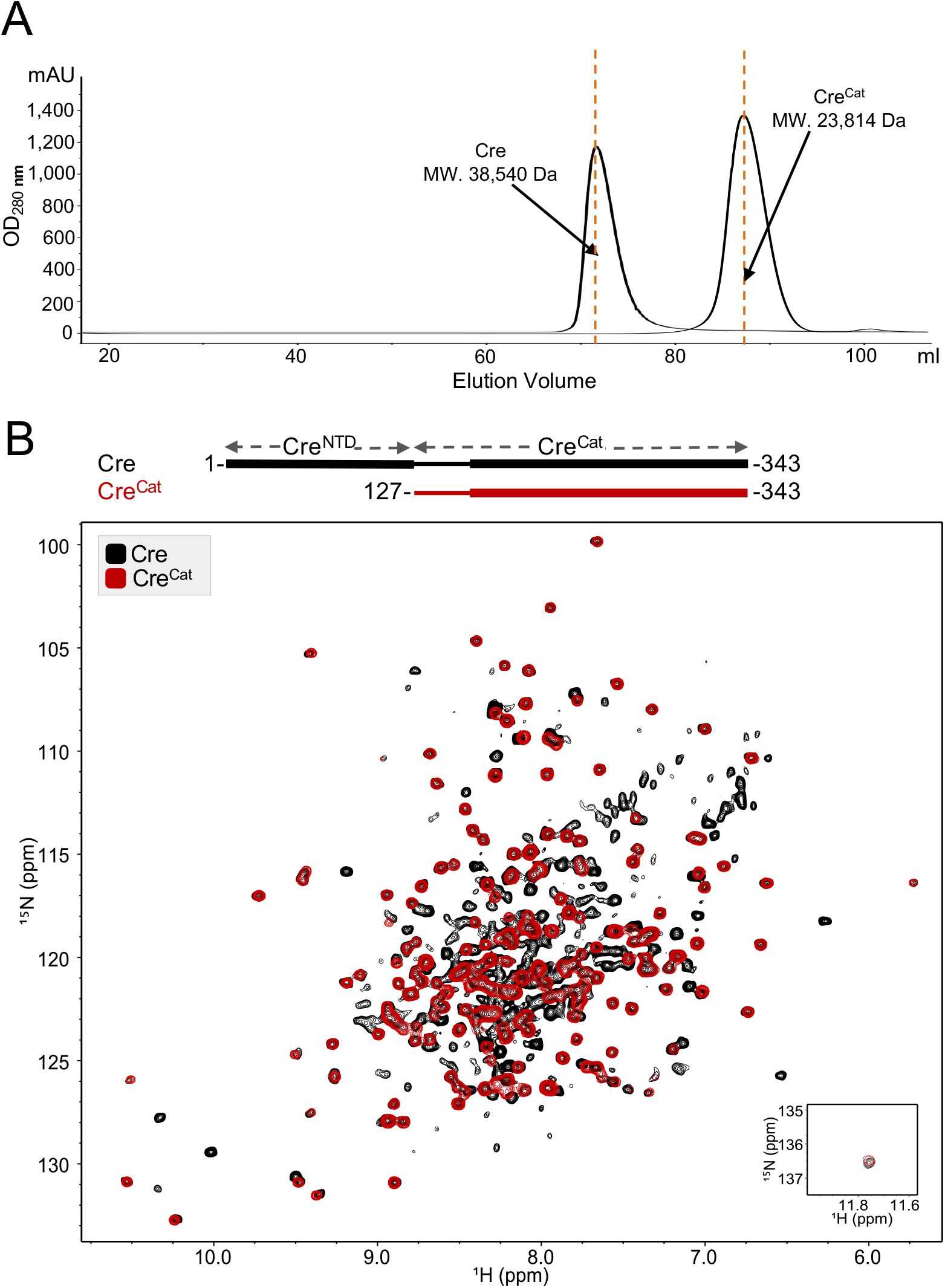
Domains of Cre fold independently in solution. *(A)* Comparing size exclusion chromatography of Cre^Cat^ domain (MW. 23,814 Da) with monomeric full-length Cre (MW. 38,540 Da) shows Cre^Cat^ is monomeric in solution. *(B)* 1H-^15^N TROSY-HSQC spectrum of Cre^Cat^ (residues 127-343; red) domain overlaid on that of full-length Cre (residues 1-343; black). Coincidence of red and black signals indicate that the domains of Cre fold independently in solution.

**Figure S2.**
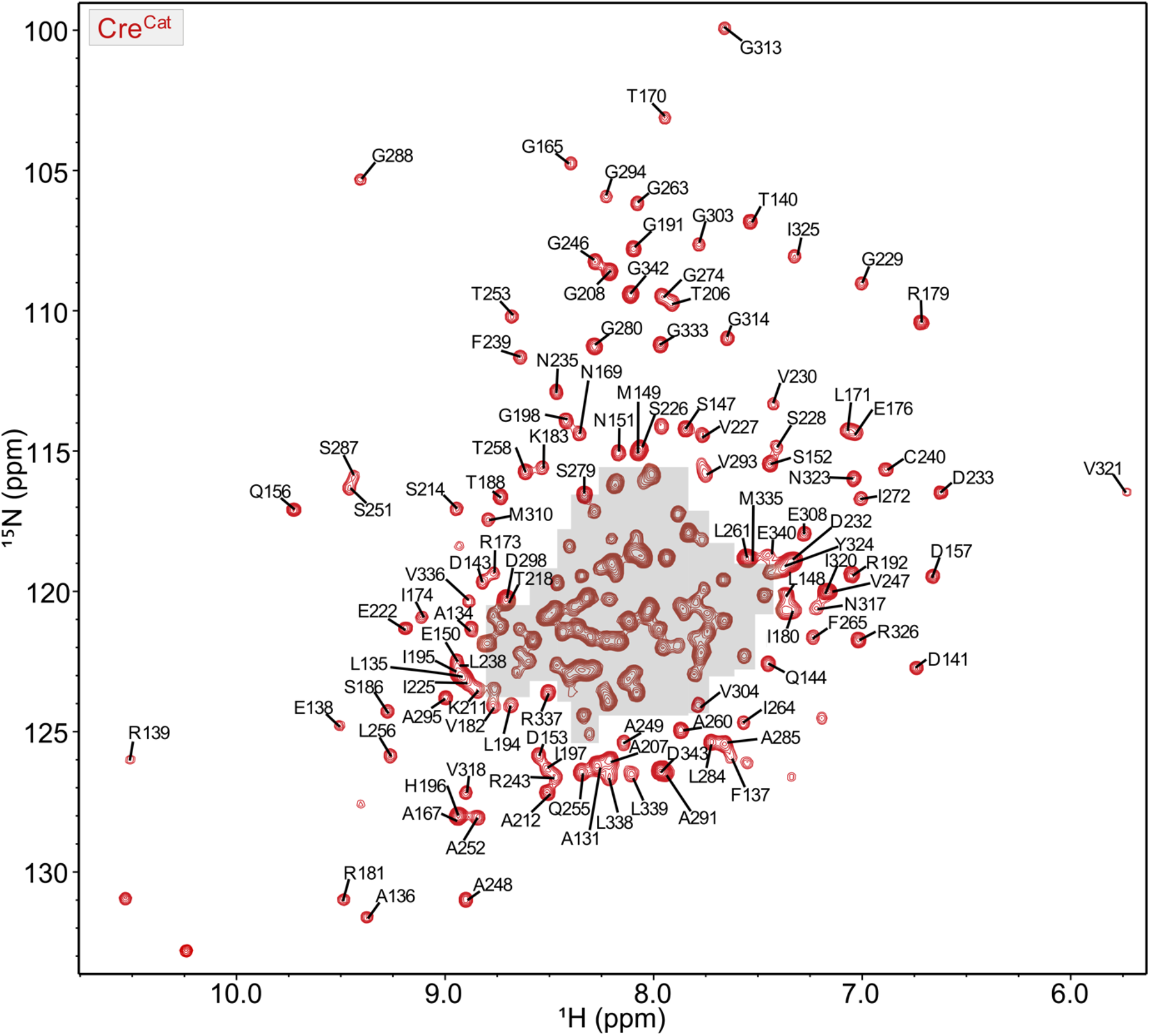
^1^H-^15^N TROSY-HSQC spectrum of Cre^Cat^: Amide chemical shift assignments are indicated; those for the crowded region highlighted in gray are shown in Fig. S3.

**Figure S3.**
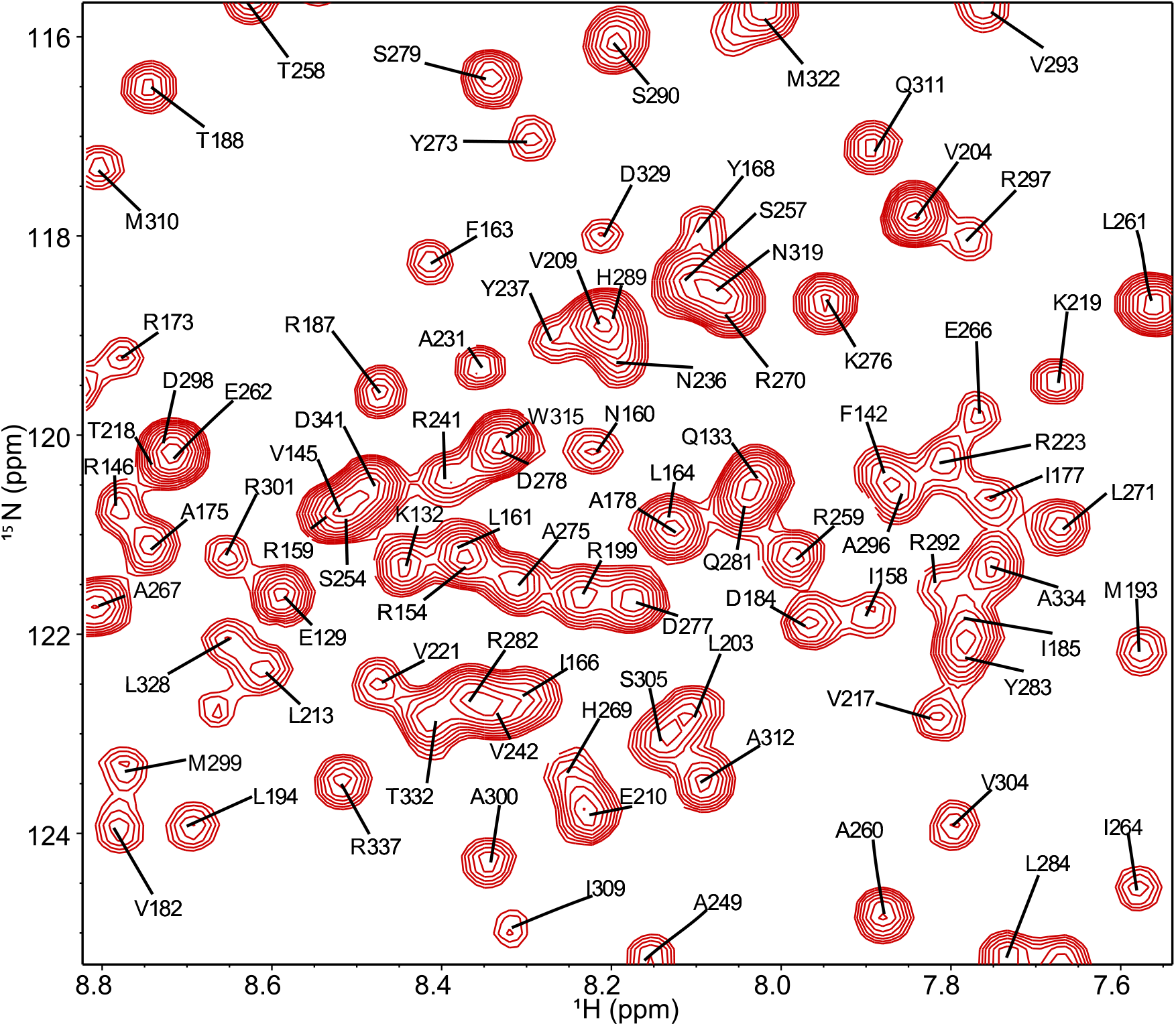
^1^H-^15^N TROSY-HSQC spectrum of Cre^Cat^ showing amide assignments from the crowded region highlighted in Fig. S2 in gray.

**Figure S4.**
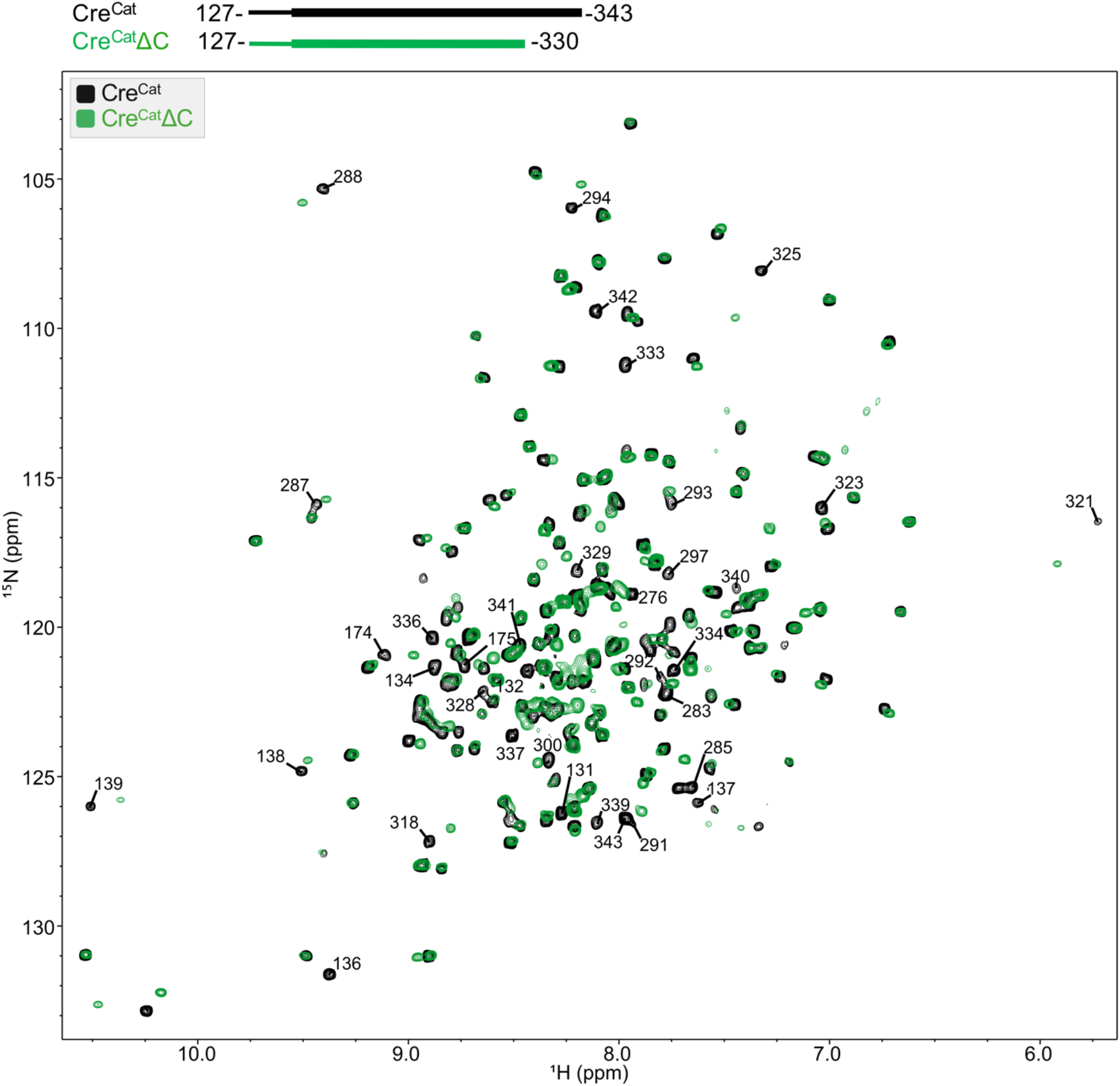
Truncation of C-terminal region causes CSPs in resonances from Cre^Cat^ core. Overlay of ^1^H-^15^N TROSY-HSQC spectra of Cre^Cat^ (residues 127-343; black) and Cre^Cat^ ΔC (residues 127-330; green). Assignments of residues with CSPs greater than 1 SD (= 0.065 ppm) are indicated.

**Figure S5.**
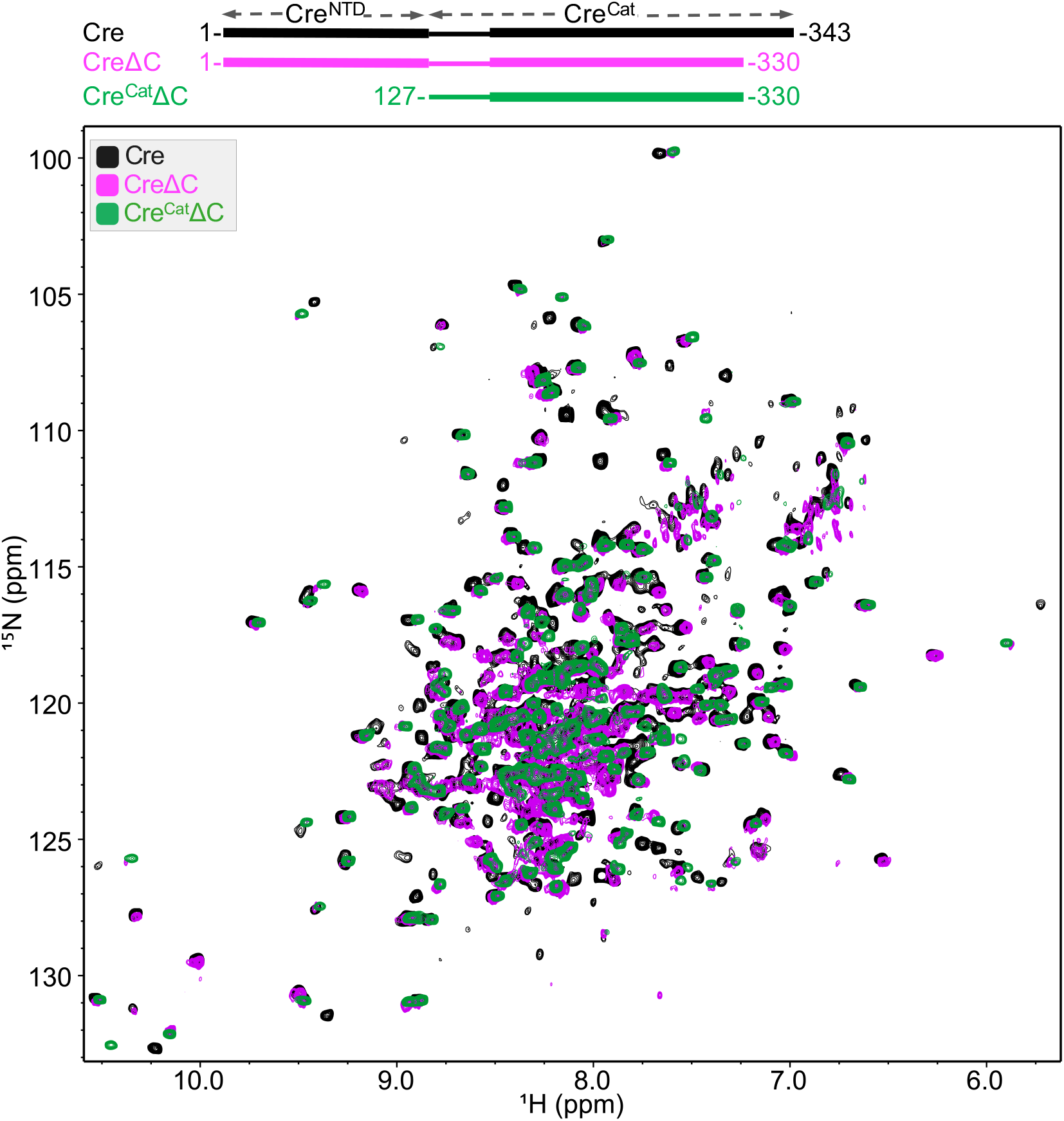
Truncation of C-terminal region does not affect the independent folding of Cre^NTD^ domain. Overlay of ^1^H-^15^N TROSY-HSQC spectrum of full-length Cre (residues 1-343; black), CreΔC (residues 1-330; magenta) and Cre^Cat^ΔC (residues 127-330; green). CSPs upon truncation of C-terminal residues Δ(E331-D343) are observed only in the catalytic domain (green Cre^Cat^ΔC peaks overlay with magenta CreΔC peaks, not with those black full-length Cre peaks, indicating same CSPs upon truncation; while rest of the peaks from black spectrum that are attributed to Cre^NTD^ overlay with corresponding magenta CreΔC peaks).

**Figure S6.**
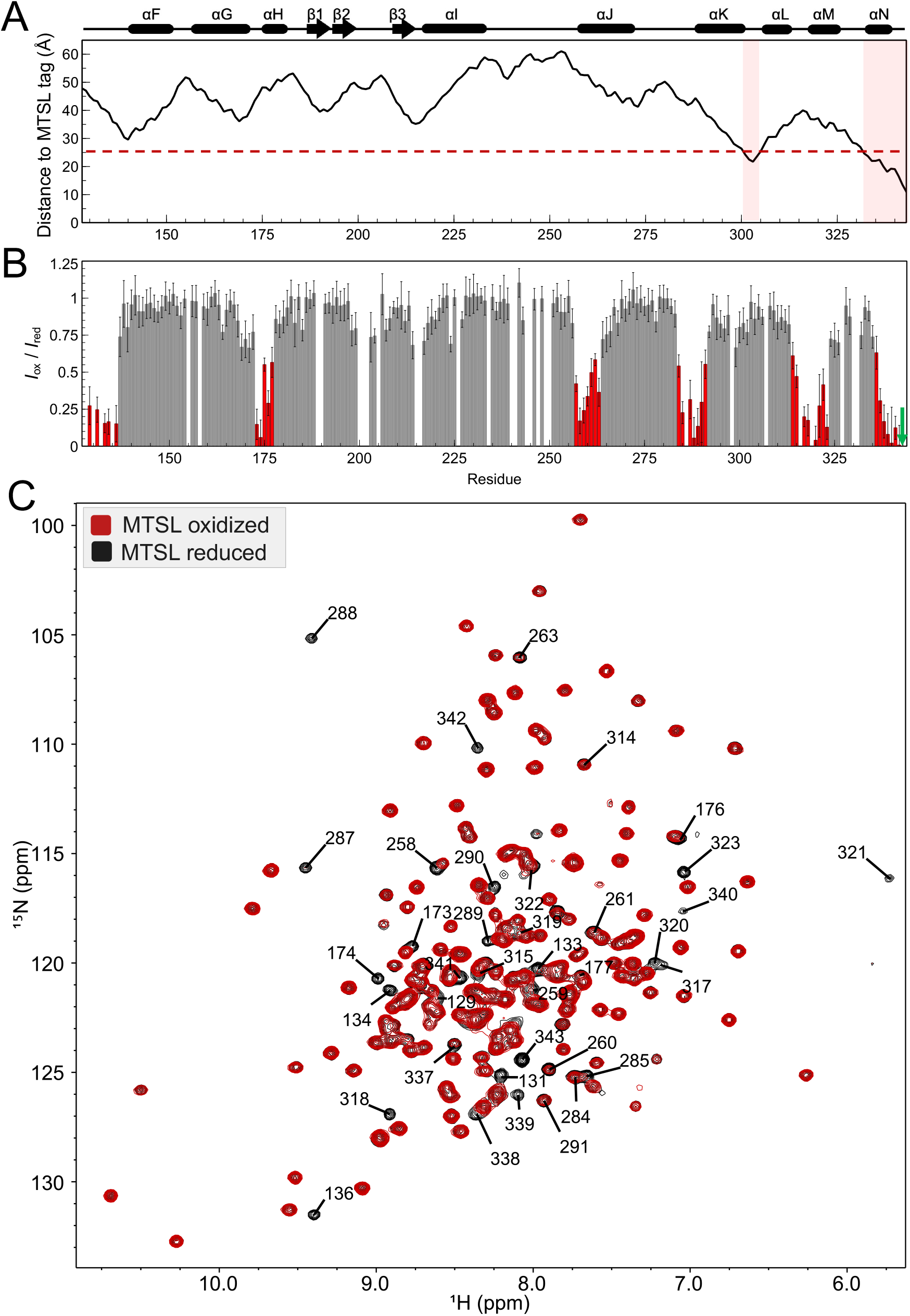
PRE-NMR experiments are consistent with *cis* docking by the C-terminus of free Cre^Cat^. *(A)* Calculated distances between backbone amide nitrogens of residues in Cre^Cat^ and MTSL-oxygen atom (distances calculated from MTSL tag manually modeled into Cre^Cat^ protomer (PDB ID 2HOI, Chain B: 127-341 and G342, C343-MTSL modeled)). Red dashed line at 25 Å represents the limit of observable PRE effects predicted from the synaptic Cre tetramer structure. Regions highlighted in pink have a calculated distance ≤ 25 Å. Secondary structure elements of Cre synaptic complex crystal structure (PDB ID 2HOI, chain B) are shown above the plot. *(B)* Observed PRE effects in MTSL tagged-free Cre^Cat^ C155A/C240A/D343C. Panel details same as Fig. 4*A*. Observed PRE effects differ significantly from the predictions shown in panel S6 *A*. *(C)* Overlay of ^1^H-^15^N TROSY-HSQC spectra of MTSL tagged-free Cre^Cat^ C155A/C240A/D343C in oxidized (paramagnetic, red) and reduced states (diamagnetic, black).

**Figure S7.**
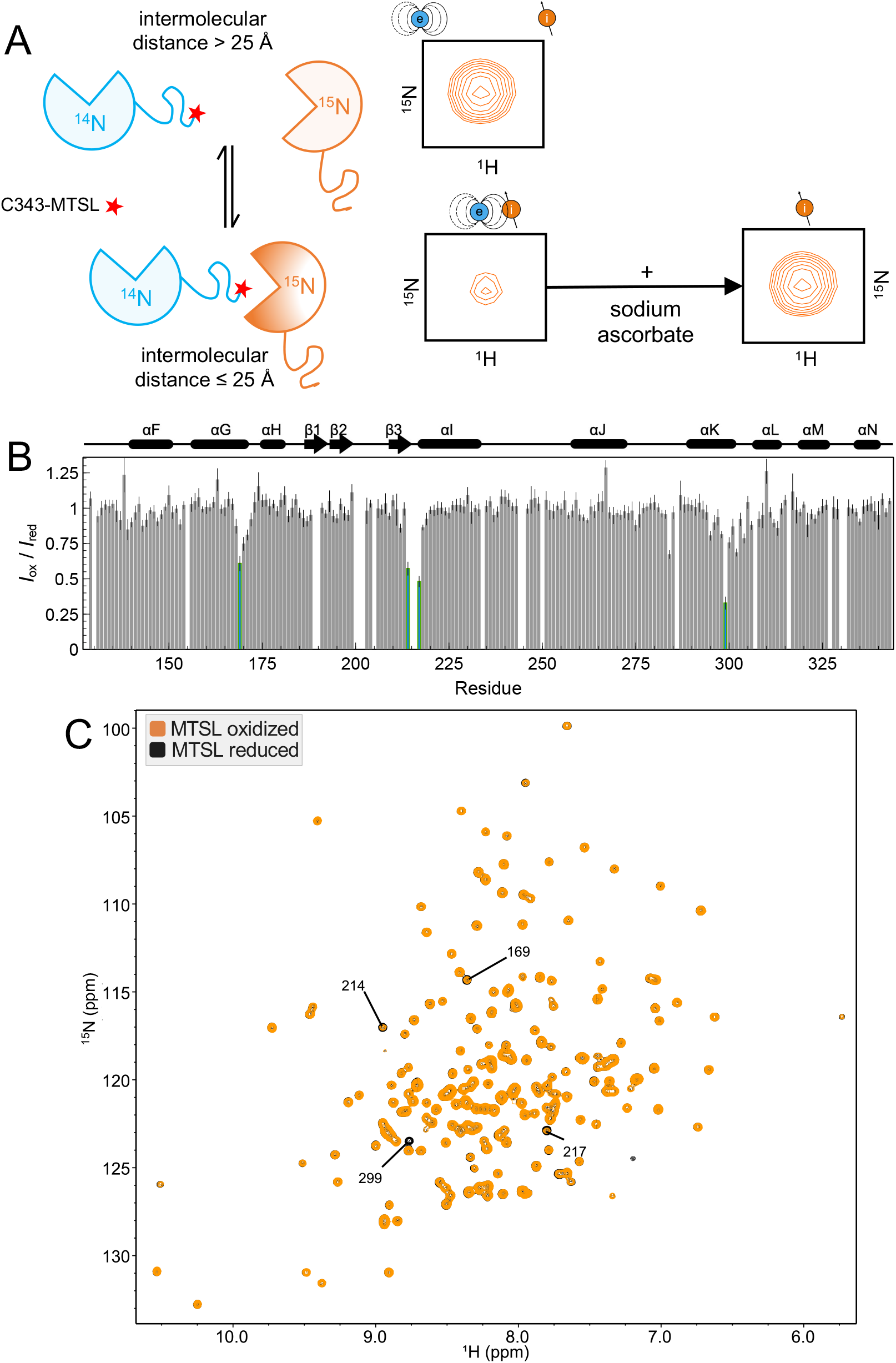
Weak intermolecular PRE effects indicate transient *trans* interactions between Cre^Cat^ molecules in the absence of DNA. *(A)* Schematic of intermolecular PRE-NMR experiment of Cre^Cat^ using MTSL-tagged unlabeled Cre^Cat^ C155A/C240A/D343C mixed with non-tagged [U-^15^N] WT Cre^Cat^ in a 1:1 molar ratio. Residues in [U-^15^N]-Cre^Cat^ at an intermolecular distance ≤ 25 Å from the C343-MTSL spin tag (star) will show a reduction in signal intensity due to free radical-induced PRE effects. Addition of excess sodium ascorbate reduces the MTSL tag to its diamagnetic state and results in normal signal intensity. *(B)* Per-residue normalized intensity ratios (*I*_ox_/*I*_red_) of the intermolecular PRE-NMR profile of free Cre^Cat^. Residues with PRE effects (*I*_ox_/*I*_red_ < 0.65) are highlighted in green. Uncertainties are propagated from the signal-to-noise ratio of individual resonances. The secondary structure elements of the Cre synaptic complex crystal structure (PDB ID 2HOI, chain B) are shown above the bar-graph. *I*_ox_/*I*_red_ ratio ~ 1 in most resonances indicates only weak transient intermolecular interactions in free Cre^Cat^. *(C)* Overlay of ^1^H-^15^N TROSY-HSQC spectra of MTSL oxidized (paramagnetic; orange) and reduced states (diamagnetic; black) for the intermolecular PRE studies of free Cre^Cat^. Only the four signals labeled showed PRE effects.

**Figure S8.**
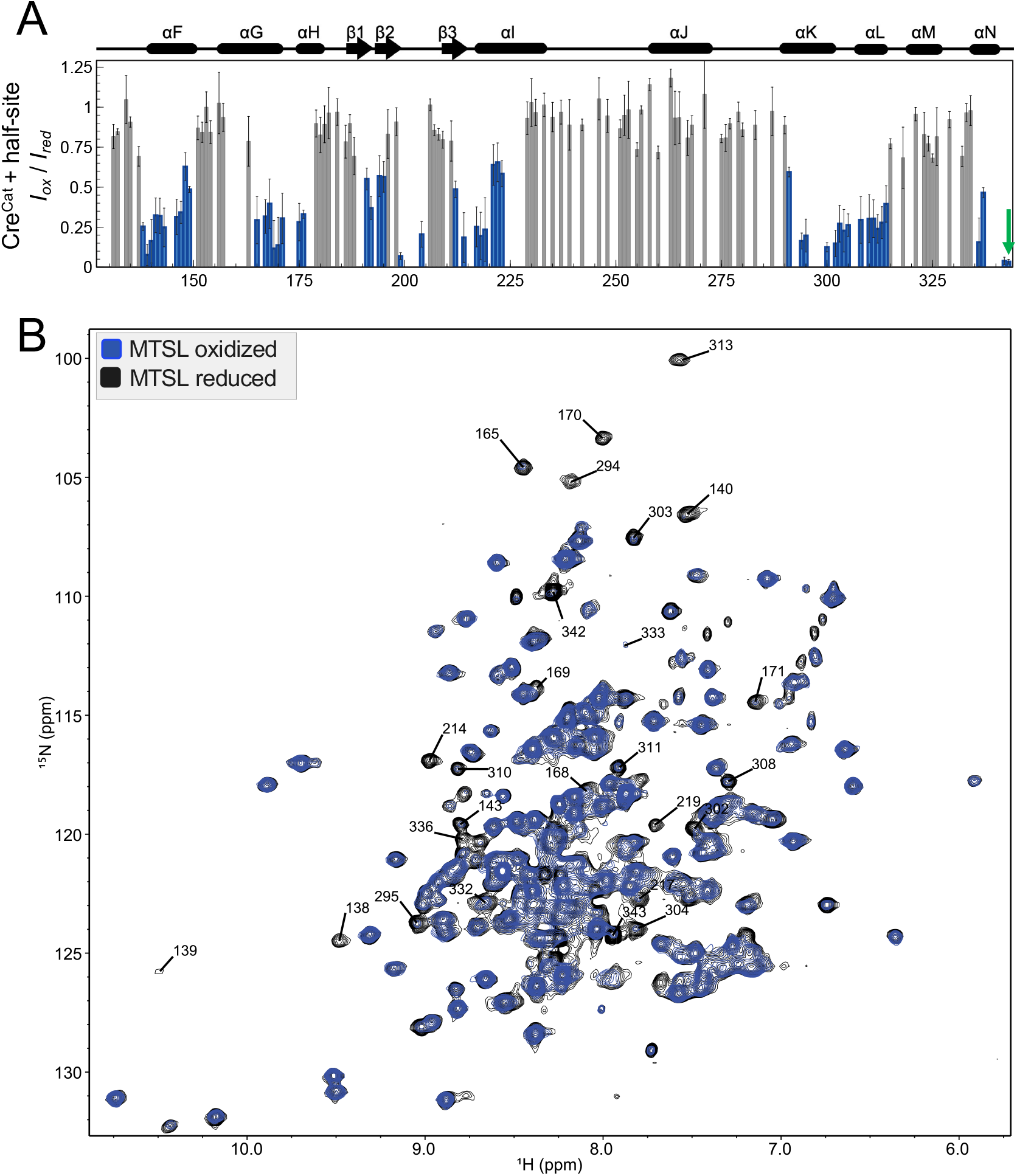
DNA bound Cre^Cat^ shows PRE effects in residues at the *trans* docking cavity. *(A)* Observed PRE effects in MTSL tagged-Cre^Cat^ C155A/C240A/D343C, in complex with *loxP* DNA half-site. Panel details same as Fig. 4*B*. *(B)* Overlay of ^1^H-^15^N TROSY-HSQC spectra of MTSL tagged-Cre^Cat^ C155A/C240A/D343C-DNA complex oxidized (paramagnetic, blue) and reduced by addition of sodium ascorbate (diamagnetic, black).

**Figure S9.**
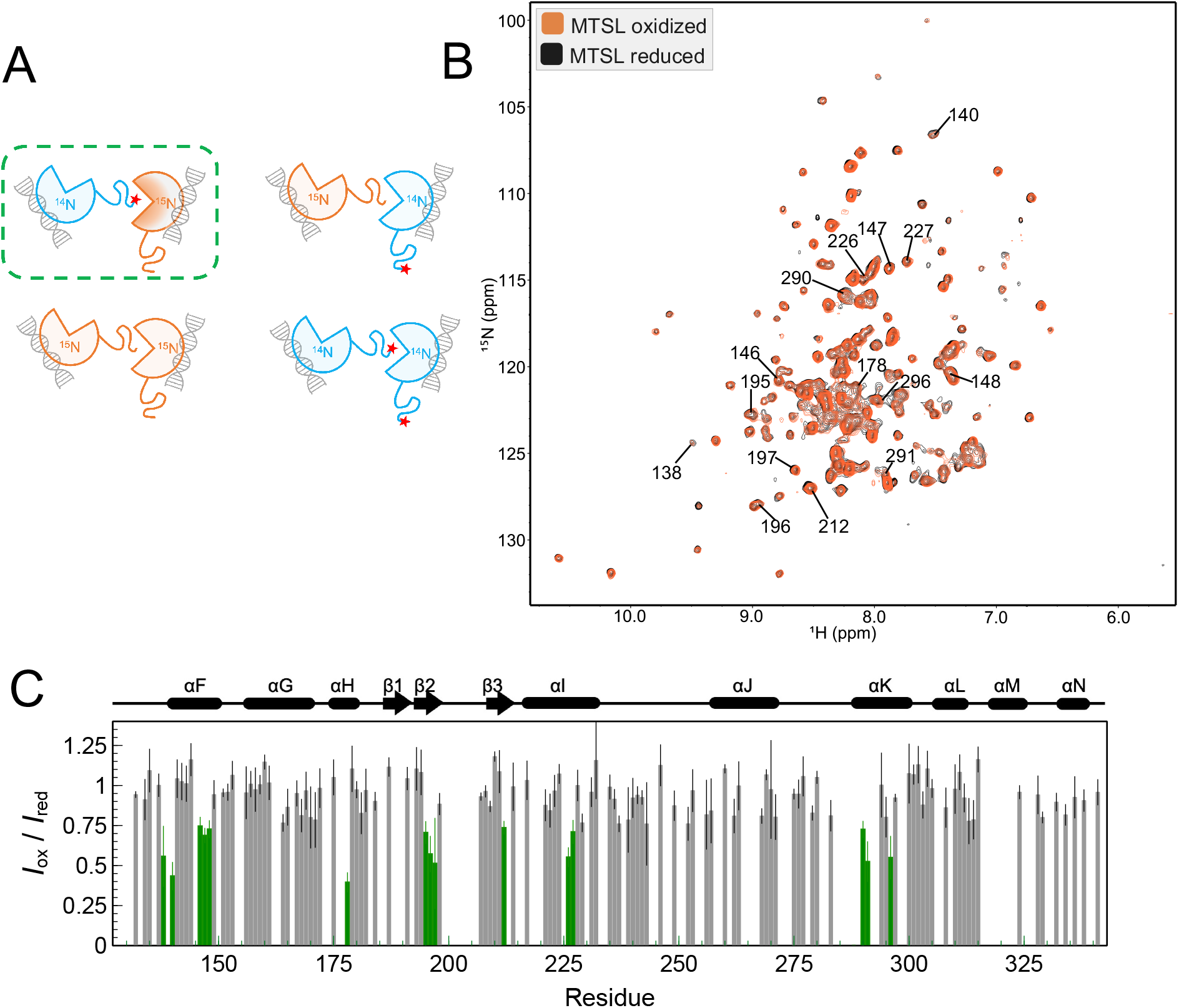
Strong Intermolecular PRE-NMR of Cre^Cat^ -DNA half-site complex is consistent with *trans* interactions via Cre C-terminus between half-site complexes. *(A)* Schematic of intermolecular PRE-NMR of Cre^Cat^-DNA half-site complexes using MTSL tagged-natural abundance Cre^Cat^ C155A/C240A/D343C and non-tagged [U-^15^N] Cre^Cat^ proteins when in complex with DNA half-site mixed in a 1:1 molar ratio. Four possible relative orientations are predicted for the interaction of protein molecules in complex with DNA according to DNA-bound tetrameric structures of Cre (PDB ID 2HOI). Of the four orientations, only the one highlighted in green would report intermolecular PRE effects. *(B)* Overlay of ^1^H-^15^N TROSY-HSQC spectra of MTSL oxidized (paramagnetic; orange) and reduced states (diamagnetic; black) in the intermolecular PRE studies of the Cre^Cat^-DNA complex. *(C)* Per-residue normalized intensity ratios (*I*_ox_/*I*_red_) of the intermolecular PRE-NMR profile of Cre^Cat^-DNA complex; *I*_ox_/*I*_red_ values < 0.7 are shown in green. Uncertainties are propagated from the signal-to-noise ratio of individual resonances. The secondary structure elements of Cre synaptic complex crystal structure (PDB ID 2HOI, chain B) are shown above the bar-graph.

**Table S1.**
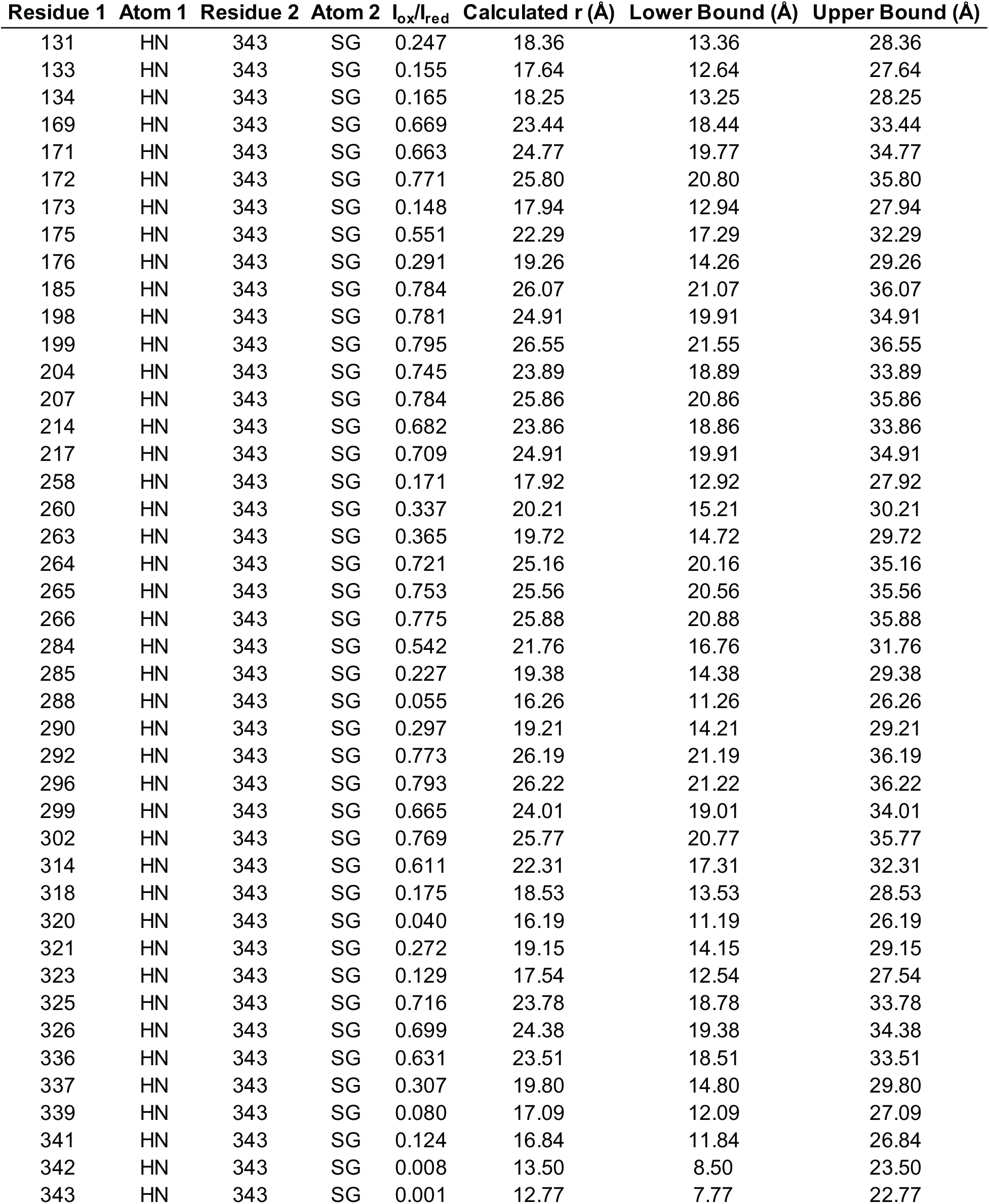
PRE-derived distance restraints for *I*_*ox*_ / *I*_*red*_ < 0.8

